# Brassinosteroid-regulated transcription factors confer epigenetic changes that repress plant immunity

**DOI:** 10.64898/2026.04.09.717176

**Authors:** Veronica E. Ramirez, Haiwei Shuai, Fang-Yu Hwu, Rashmi R. Hazarika, Chia-Nan Tao, Sera Choi, Robert S. Piecyk, Susanne I. Wudy, Michael Gigl, Johannes W. Bagnoli, Sarah Brajkovic, Pablo Albertos, Yuanyuan Liang, Andreas Keymer, Corinna Dawid, Wolfgang Enard, A. Corina Vlot, Caroline Gutjahr, Martin Parniske, Bernhard Kuster, Tobias Sieberer, Christina Ludwig, Cyril Zipfel, Jurriaan Ton, Frank Johannes, Brigitte Poppenberger

## Abstract

When organisms encounter pathogens, they rapidly activate complex defense programs to ensure survival. While these immune responses are vital, they often also incur trade-offs, such as reduced growth and development and must therefore be tightly controlled ^1,2^. In this study, we reveal that the steroid hormones brassinosteroids (BRs) contribute to this control in *Arabidopsis thaliana* by repressing immunity-related genes. We provide evidence that the BR-regulated basic helix-loop-helix (bHLH) transcription factor CESTA (CES), along with its homologs BR ENHANCED EXPRESSION (BEE)1-3, mediate DNA methylation changes at transposable element (TE)-rich loci containing nucleotide-binding leucine-rich-repeat (NLR)-type receptor genes, including *SUPPRESSOR OF NPR1-1 CONSTITUTIVE 1* (*SNC1*). These CES-induced methylation changes correlate with altered splicing of *SNC1* pre-mRNA, a process that requires the BR receptor BRASSINOSTEROID INSENSITIVE 1 (BRI1). In support, we show that CES associates with components of the chromatin remodeling and splicing machinery. Together, our findings reveal a previously unrecognized BR-induced mechanism that modulates the epigenetic and post transcriptional regulation of immune genes, enabling plants to prioritize growth over defense.

**Significance statement:** Steroid hormones are powerful regulators of growth but also act as potent suppressors of immunity, with well-established clinical applications, for example in treating autoimmune diseases in humans. In plants, the steroid hormones brassinosteroids (BRs) exert similar effects. Here, we uncover that BR-mediated immune suppression involves DNA methylation changes and alternative splicing of a subset of immune receptor genes, that govern resistance responses mediated by the plant hormone salicylic acid. We provide evidence for a function of specific bHLH transcription factors in this process that link BR signaling to chromatin remodelling and RNA processing. Thereby our study reveals a novel mode of steroid hormone activity in immune repression, which involves epigenetic and posttranscriptional adjustment of immune receptor function.

## Introduction

The immune system of plants consists of different layers that recognize microbial invaders at multiple steps of the infection process to induce mechanical and chemical defenses against such perpetrators. Pathogen perception occurs through plasma-membrane-bound pattern recognition receptors (PRRs) that detect pathogen-associated molecular patterns (PAMPs) and intracellular receptors such as nucleotide-binding leucine-rich-repeat receptors (NLRs) that perceive effectors released by pathogens and induce pattern-triggered (PTI) and effector-triggered immunity (ETI). The downstream reactions that then follow are multilayered and include increased ion influx and oxidative bursts, controlled cell death, and the activation of immune-responsive genes via signaling cascades mediated by hormones, such as salicylic acid (SA) ^3,4^.

Immune responses are metabolically costly and often yield trade-offs, such as inflammation related diseases in humans ^5^ or growth inhibition in plants; this is particularly evident in autoimmune mutants that display severe dwarfism ^6^. To minimize these costs, they are closely regulated and one key mechanism implicated is DNA methylation, an epigenetic mark that silences potentially harmful genetic elements and modulates gene expression ^7,8^. DNA methylation can occur in promoters, which affects transcription, or in gene bodies, which influences splicing and other RNA processing means ^9,10^. In plants, it is established *de novo* by the enzyme DOMAINS REARRANGED METHYLTRANSFERASE 2 (DRM2) in CG, CHH, or CHG sequence contexts (where H = A, C, or T) via the RNA-directed DNA methylation (RdDM) pathway, is maintained by specific maintenance methyltransferases, and can be actively removed by DNA glycosylases ^11^.

While it is known that DNA methylation functions in immunity, the mechanisms that drive locus specific changes and their specific effects on gene expression remain largely elusive. Emerging evidence suggests that hormones may direct these and other related epigenetic modifications and BRs are candidates, as they can suppress biotic stress resistance ^3,12^. BRs are structurally similar to mammalian steroid hormones and signal through a phosphorylation cascade initiated at the cell surface by the receptor kinase BRASSINOSTEROID INSENSITIVE 1 (BRI1) and its coreceptor BRI1-ASSOCIATED KINASE 1 (BAK1) ^13^. Notably, BAK1 also serves as co-receptor for the PRR FLAGELLIN SENSING 2 (FLS2), which activates immune signaling and this shared role of BAK1 in BR-mediated growth regulation and immune responses contributes to growth repression when immune responses are activated ^14^.

In addition to receptor activity, downstream components of BR signaling also contribute to immune suppression. This has been demonstrated for the BR-regulated transcription factors (TFs) BRASSINAZOLE RESISTANT 1 (BZR1) and HOMOLOGUE OF BEE2 INTERACTING WITH IBH1 (HBI1). Overexpression of BZR1 and HBI1 reduced resistance to *Pseudomonas syringae* pv. *tomato*, an effect associated with suppressed reactive oxygen species (ROS) bursts in response to both fungal and bacterial elicitors, as well as the downregulation of SA-responsive genes ^15-18^.

HBI1 is a bHLH TF and a close homolog of CETSA (CES) and the BR ENHANCED EXPRESSION (BEE) 1-3 proteins, which are induced by BRs at both transcriptional and posttranslational levels ^19,20^. Upon activation of BR signaling, CES and the BEEs accumulate in distinct subnuclear compartments. In case of CES, this nuclear compartmentalization depends on SUMOylation ^21^, a post-translational modification known to promote the formation of nuclear bodies and influence chromatin dynamics, including changes in DNA methylation, in both animals and plants ^22,23^.

## Results

### CES Represses *Hpa* Resistance

CES functions as a positive regulator of several subsets of BR-induced genes ^20,24,25^. However, microarray analysis of the dominant, gain-of-function (*gof*) *ces-D* mutant, an activation-tagged line, where the integration of a 4x35S enhancer element in the promoter leads to a strong, constitutive induction of CES, had revealed that CES can also repress gene expression ^20^. Among the *ces-D*-repressed genes were 20 NLRs, several of which exhibited a 50-100-fold reduction (*SI Appendix*, Table S1); they included seven members of the *RESISTANCE AGAINST PERONOSPORA PARASITICA* 5 (*RPP5)* gene cluster, which comprises eight closely related NLRs and several TEs in the *A. thaliana* Columbia-0 (Col-0) ecotype ^26,27^. The RPP5-cluster contains *RPP4* and *SUPPRESSOR OF NPR1-1 CONSTITUTIVE 1* (*SNC1*), encoding NLRs that mediate resistance against *Hyaloperonospora arabidopsidis* (*Hpa)*, the causal agent of downy mildew in *A. thaliana* ^28,29^.

To assess whether CES influences *Hpa* resistance, infection assays were performed using the virulent *Hpa* isolate Noco2. Alongside *ces-D*, the CES over-expression line *35S:CES-YFP* (*CESoe*)^20^, and two loss-of-function (*lof*) mutants - a *ces bee1,3* triple mutant (*ces-tM*) and a *ces bee1,2,3* quadruple mutant (*ces-qM*) ^25^-were analyzed in comparison to wild-type (wt) plants. The results of oomycete colonization assays showed that resistance, evidenced by reduced colonization, was significantly increased in the *ces lof* mutants and significantly decreased in *cesD* as compared to wt (Fig. 1*A*). The *CESoe* plants showed a tendency towards increased *Hpa* colonization, although the difference was not statistically significant. In confirmation, spore and sporangiophore formation were significantly reduced in the two *ces lof* mutants and increased in the two *gof* lines, although the *gof* phenotypes were again more variable (Fig. 1*B*).

**Fig. 1.**
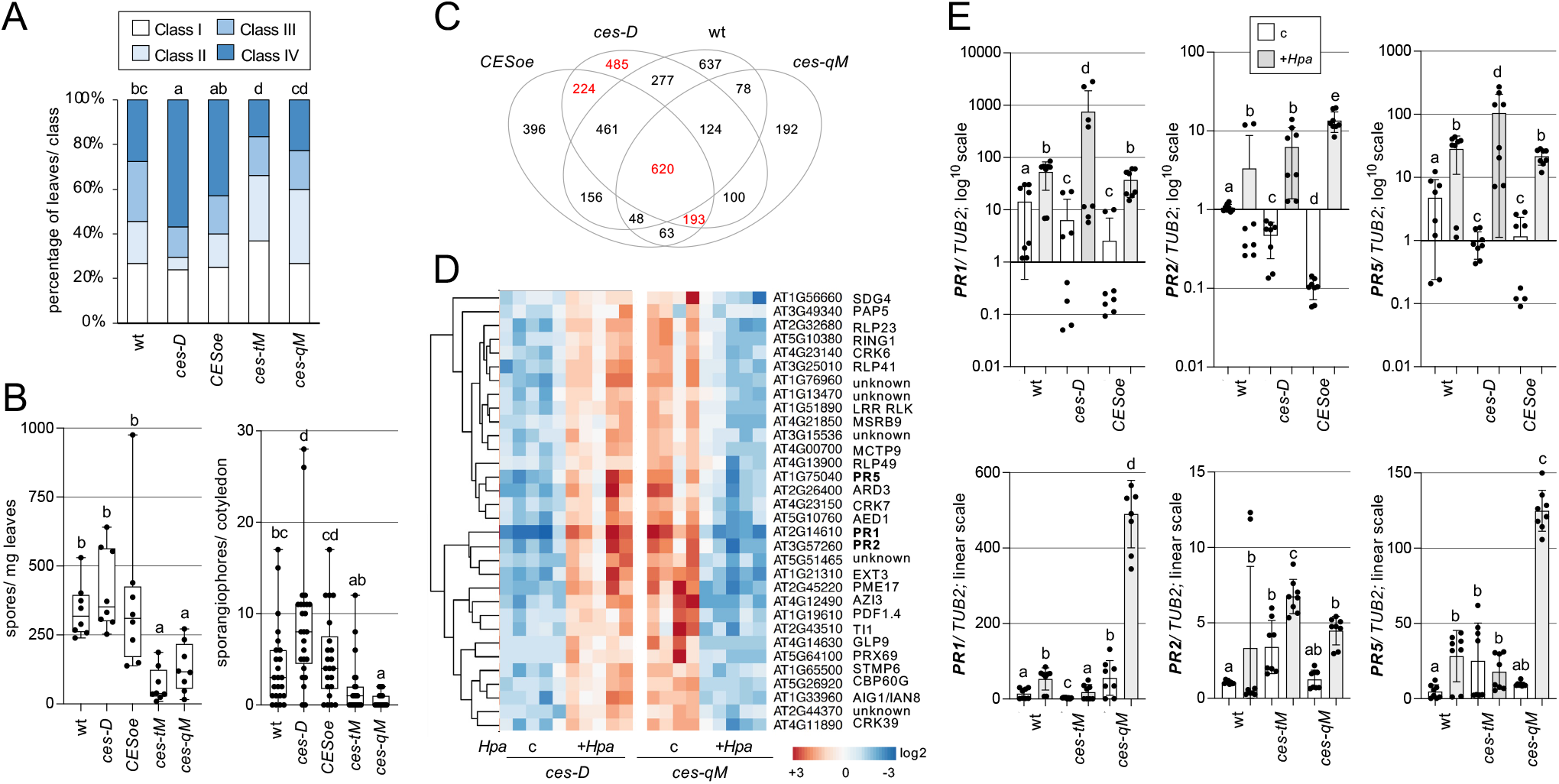
CES represses *Hpa* resistance responses. (*A, B*) *Hpa* colonization and growth in *ces* mutants. Two-week-old seedlings were spray-inoculated with *Hpa* Noco2. a, Microscopic analysis of trypan blue-stained leaves at 6 dpi. *Hpa* colonization was scored and categorized into four infection classes; data represent the percentage of leaves per class (n = approx. 100 leaves from 20-25 plants per genotype), significance determined with pairwise Fisher’s exact tests. b, Quantification of spores (per mg leaf fresh weight) and sporangiophores (per cotyledon) at 5 dpi (n = 8-15 and 24 respectively) significance determined with two-sided Mann–Whitney U-tests. (*C*) Venn diagram showing DEGs identified by RNA-seq in *ces* mutants upon *Hpa* infection. (*D*) Heat map of expression changes in *ces-D* and *ces-qM* mutants. A subset of highly regulated genes from the 193 DEG group shown in *C* is displayed. (*E*) Absolute expression levels of *PR1, PR2*, and *PR5* in *ces gof* (upper row, log scale) and *ces lof* (lower row, linear scale) mutants relative to wt, with or without *Hpa* Noco2 infection, as determined by qPCR. Data represent mean ± SD from four biological replicates, each measured in triplicate and normalized to *TUB2*.

### CES is Required for Balanced SA Responses

Plants with constitutively active NLRs, such as the dominant *snc1* mutant ^28^, display autoimmune phenotypes characterized by strongly increased SA levels (*SI Appendix*, Fig. S1*A*) and severely stunted growth ^30^. In contrast, the *ces lof* lines did not show highly elevated SA levels (*SI Appendix*, Fig. S1*B*), consistent with the absence of an autoimmune dwarf phenotype in these plants ^24^, and indicating that CES does not constitutively activate SA responses.

To explore the molecular basis by which CES influences *Hpa* resistance, we conducted a RNA sequencing (RNA-seq) analysis of gene expression differences between wt and the four *ces* lines, both under basal conditions and in response to infection. Material of plants, either spray inoculated with the *Hpa* isolate Noco2 or with water as a control, was processed using the primeseq protocol^31^, and differential gene expression was analyzed with DESeq2 to quantify log2-fold changes. Cluster analysis of the RNA-seq data revealed tight clustering of all wt and *gof* samples, indicating high consistency across replicates. In contrast, one replicate each of treated and control group *ces-qM* as well as all *ces-tM* samples failed to cluster (*SI Appendix*, Fig. S2*A*) and these were therefore excluded from further analysis.

A comparison between *Hpa*-treated and control samples within each genotype identified 620 differentially expressed genes (DEGs) that responded to *Hpa* in all *ces* mutant and wt plants (FDR < 0.05). These 620 DEGs clustered into two major groups: cluster 1, genes upregulated, and cluster 2, genes downregulated following *Hpa* infection in wt and the two *gofs* (*SI Appendix*, Table S2). Importantly, this regulation was stronger in the *gofs* than in wt and was reverted in *ces-qM*, providing evidence for a role of CES in controlling large aspects of *Hpa*-induced transcriptome changes. Moreover, the analysis showed that cluster 1 genes were constitutively activated and less induced following infection in *ces-qM*, compared to wt and the *ces gofs*, speaking for CES taking part, not only in the induction and repression of these genes in response to *Hpa*, but also in the control of their basal levels in the absence of a pathogen.

The majority of genes in Cluster 1 had a biotic stress context, confirmed through PANTHER GO biological process annotation (*SI Appendix*, Fig. 2*B*). Moreover, a PANTHER GO functional annotation revealed significant enrichment of oxidoreductase activity, peroxidase activity, and oxidoreductase activity acting on peroxide as an acceptor (*SI Appendix*, Fig. 2*C*). A comparison with a list of 171 identified markers of ROS-responses ^32^ revealed 43 shared genes (*SI Appendix*, Table S3), including the TFs *ZINC FINGER OF ARABIDOPSIS THALIANA 10 (ZAT10), ZAT12*, the WRKY TFs *WRKY6, WRKY33, WRKY40*, and *WRKY75*, as well as *FAD-LINKED OXIDOREDUCTASE 1* (*FOX1*), providing evidence that CES is required for the repression of ROS responses.

**Fig. 2.**
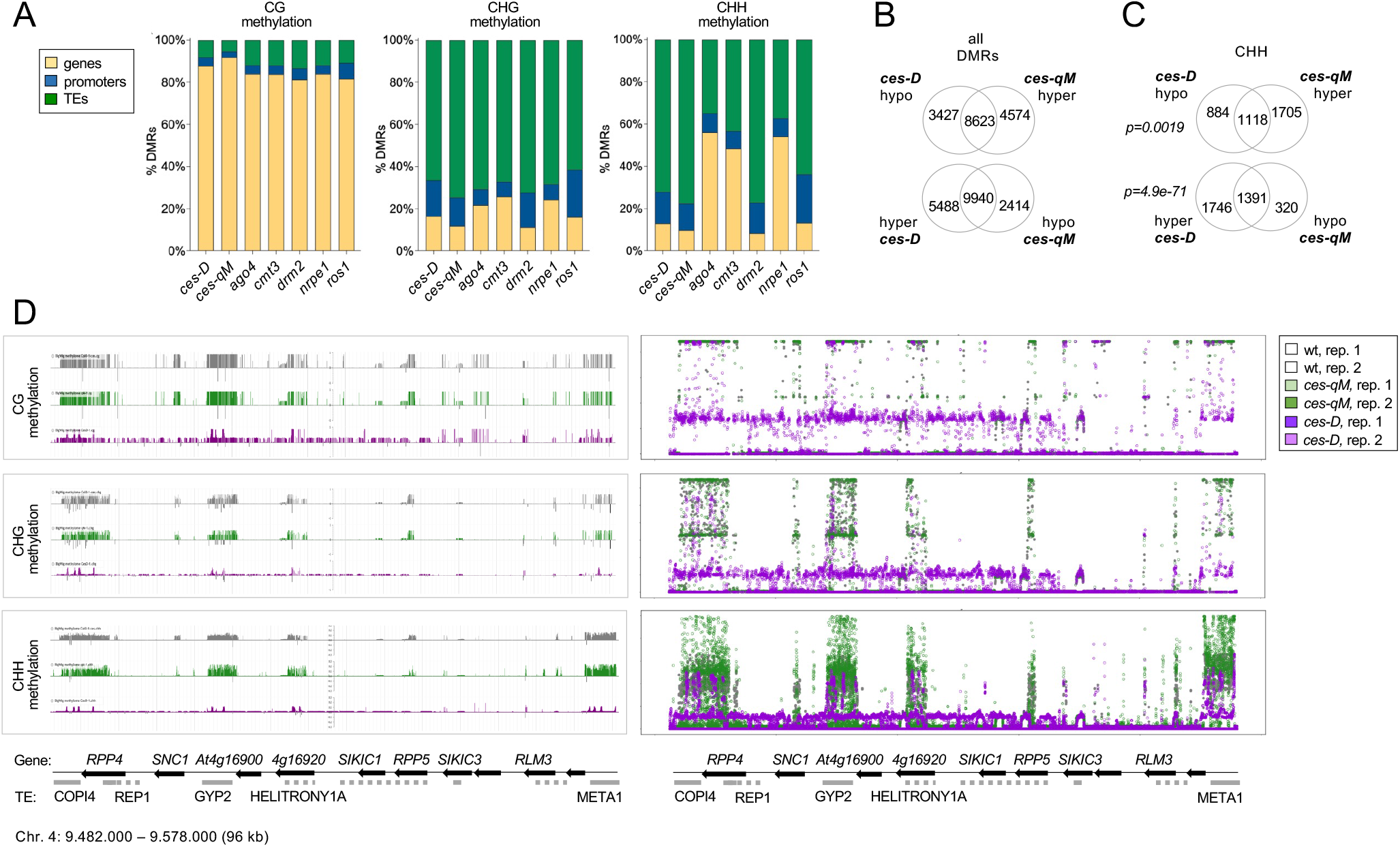
CES globally alters DNA methylation, with the *RPP5* gene cluster region being significantly affected. (*A*) Global distribution of DMRs in *ces* mutants relative to wt, categorized by CG, CHG, and CHH sequence contexts. DMRs were identified using METHimpute; profiles are compared to known DNA methylation and demethylation mutants. (*B, C*) Venn diagrams showing overlap of hypo- and hyper-methylated DMRs between *ces-qM* and *ces-D* mutants at the genome-wide level (*B*) and restricted to the CHH context (*C*). (*D*) Visualization of methylation changes at the *RPP5* gene cluster. Epigenome browser views (JBrowse, left) and bubble plots (right) display strand-specific methylation in CG, CHG, and CHH contexts for wt (grey), *ces-qM* (green), and *ces-D* (purple). Y-axes indicate methylation proportion per context; scales are automatically adjusted per dataset. Gene and TE annotations for the region are shown below.

193 DEGs exhibited altered responsiveness to *Hpa* in CES *lof* and *gof* mutants only, evidencing a stronger regulation than in wt (Fig. 1*C*; *SI Appendix*, Table S4). Among these mutant-specific DEGs, the majority were basely repressed and showed enhanced responsiveness to *Hpa* in the two *gof* lines. In contrast, they were constitutively upregulated in *ces-qM*, displaying weaker *Hpa-* induced expression as compared to *ces-D* (*SI Appendix*, Table S4). Notably, this subset included many SA-responsive genes, such as the *PATHOGENESIS-RELATED* (*PR*) genes *PR1, PR2*, and *PR5*, as well as the SA signaling regulators *PHYTOALEXIN DEFICIENT 4* (*PAD4*) and *ENHANCED DISEASE SUSCEPTIBILITY 5* (*EDS5*) (Fig. 1*D, SI Appendix*, Table S4). Moreover, the *ces* mutant-specific DEGs included several receptor-encoding genes, such as the *CYSTEINE-RICH RECEPTOR-LIKE PROTEIN KINASES CRK4, CRK6, CRK7, CRK14, CRK28*, and *CRK36*, the *RECEPTOR-LIKE PROTEINS RLP23* and *RLP49*, and receptor kinases such as *BAK1-INTERACTING RECEPTOR-LIKE KINASE 1 (BIR1*) and *SUPPRESSOR OF BIR 1 (SOBIR1*).

To validate the RNA-seq results, quantitative real-time PCR (qPCR) was performed on *PR1, PR2*, and *PR5*, which are well-established markers of SA responses. The qPCR data confirmed a reduced basal expression of these genes in the two *gof* mutants, while *cesqM* exhibited elevated basal levels. Additionally, all three genes showed a stronger induction upon *Hpa* infection in the *ces* mutants compared to wt (Fig. 1*E*). Expression patterns in *cestM* were more variable, aligning with previous evidence of functional redundancy among the BEE branch of bHLH TFs ^17,19^, which may contribute to the less stable outcomes in the lower-order mutant setting. Interestingly, a member of this subfamily, HBI1, which, like CES, can activate BR biosynthesis ^17^, does not appear to operate in this specific regulatory branch within BR-immunity crosstalk, since *HBI1* over-expression neither altered *SNC1* splicing nor *Hpa* Noco2 resistance (*SI Appendix*, Fig. S3A-C). In support, a comparison of overlapping targets between exclusively *ces-D*-containing *Hpa*-regulated subgroups and *HBI1oe* RNA seq-derived DEGs ^33^ showed largely opposite transcriptome changes (*SI Appendix*, SFig. 3*D* and Table S5), indicating distinct rather than redundant roles of CES and HBI1.

A mass-spectrometry-based proteome analysis was then utilized to determine if CES-conferred transcriptome changes are translated to the proteome. When comparing the *ces-qM* mutant to wt without and with *Hpa* treatment, 258 major protein IDs were identified as being significantly altered in abundance in *ces-qM* (FDR < 0.01; fold change ≥ 2); of those, 205 were increased in basal conditions and 140 were increased following infection, with an overlap of 87 proteins that were elevated in *ces-qM* in both settings (*SI Appendix*, Fig. S4*A, B* and Table S6). Among the 258 differentially abundant proteins, 41 were encoded by genes that were part of the 193 mutant-specific DEGs, a highly significant overlap (χ^2^ test, p=4.3e-40) and these included PR1, PR2, PR5, PAD4, CRK6, CRK7, CRK14, and RLP23 (*SI Appendix*, Table S7). Thus, the proteome analysis confirmed the transcriptome changes in SA responses.

This included support for a role of CES in repressing ROS responses since in the *ces-qM* mutant several key ROS-related proteins were elevated compared to wt, including ALPHADIOXYGENASE 1 (DOX1) and FOX1, as well as multiple biosynthetic enzymes, such as the peroxidases PER50, PER51, PRXIID, PRX16, PRX37, PRX62, PRX69, PRX71, and TPX2 (*SI Appendix*, Table S6). Consistently, in ROS burst assays, using the bacterial elicitors flg22 or elf18, ROS production was reduced in the two *gof* lines and slightly increased in *ces-qM* when elf18 was utilized (*SI Appendix*, Fig. S4*C*).

In summary, these findings indicate that CES functions as a negative regulator of a wide array of immune receptors and downstream signaling components. It plays a central role in repressing basal immunity and enabling the controlled activation of immune responses upon infection.

### CES affects DNA methylation

Because the *ces-D* mutant exhibited widespread repression of entire NLR clusters, and DNA methylation, together with other epigenetic modifications, is known to regulate NLR expression ^34^, we hypothesized that altered DNA methylation patterns may underlie aspects of the *ces* mutant phenotypes. To test this, whole-genome bisulfite sequencing (WGBS) was performed on DNA extracted from non-infected *ces-D, ces-qM*, and wt plants.

Differentially methylated regions (DMRs) in the *ces-D and ces-qM* mutants relative to wt were identified using jDMR ^35^. Detected DMRs were categorized by sequence context, CG, CHG, or CHH, and by genomic location, in promoters, gene bodies, or TEs (*SI Appendix*, Table S8). This created distinct methylation profiles for both *ces* mutants, each showing widespread and analogous differences from wt (Fig. 2*A*).

To infer which DNA (de)methylation pathways might be affected by CES, publicly available WGBS datasets from the DNA (de)methylation mutants *ago4-5, cmt3-11, drm2-3, nrpe1-11*, and *ros1-4*, were analyzed using the same bioinformatics pipeline. The DMR profiles of the *ces* mutants most closely resembled those of the *drm2* and *ros1* (Fig. 2*A*), suggesting that CES affects loci targeted by these components of DNA methylation and demethylation pathways.

Direct comparison of *ces-D* and *ces-qM* revealed that many DMRs occurred at the same genomic regions but in opposite methylation states, with regions that were hypo-methylated in *ces-D* being hyper-methylated in *ces-qM*, and vice versa, making them candidates as primary CES targets (Fig. 2*B*). This effect was especially pronounced in the CHH context, where a substantial overlap of DMRs was observed (Fig. 2*C*).

To explore a potential CES-mediated regulation of NLRs specifically, DMRs for *RPP5-, RPP2A/B-, RPP7*-, and *WHITE RUST RESISTANCE 4 (WRR4)*-containing chromatin regions were extracted and summarized in tables (*SI Appendix*, Tables S9-12). In addition, methylation patterns were visualized using genome track views generated with the Plant Epigenome Browser, alongside custom bubble charts depicting each methylation event as a color-coded marker for a more detailed resolution. This exposed pronounced differences in methylation between *ces* mutants and wt with the most prominent changes in CHH methylation within TE-rich areas.

Differential methylation was very clear in the *RPP2A/B-, RPP7-*, and *WRR4*-containing regions (*SI Appendix*, Tables S10-12 and Fig. S5), but was especially striking in the *RPP5* cluster, where TEs showed CHH hyper-methylation in *ces-qM* and CHH hypo-methylation in *ces-D* (Fig. 2*D*; *SI Appendix*, Table S9). Notably, these methylation changes extended beyond the TEs themselves into adjacent genes. In addition, differential methylation was observed within intron-containing areas of *RPP4, SNC1, At4g16900*, and *RPP5* (named *SIDEKICK 2, SIKIC 2*, in Col-0), many of which, though not all, are predicted to comprise TEs (Fig. 2*D*; *SI Appendix*, Table S9). These findings reveal a role for CES in modulating methylation at immune receptor-containing loci, particularly in CHH methylated TE-enriched regions.

Hypo-methylation can activate TEs, and as a compensatory mechanism, adjacent genes can undergo methylation via RdDM, which occurs primarily in the CG context ^36^. Consistent with this, genes within the *RPP5* cluster exhibited CG hyper-methylation in *ces-D* and CG hypomethylation in *ces-qM* (Fig. 2*D*; *SI Appendix*, Table S9). This affected several *NLRs* in the *RPP5* cluster including *SIKIC1* and *SIKIC3*, which are required for SNC1-mediated immunity ^37^. The pattern of inverse CG methylation in TE-adjacent loci extended beyond the *RPP5* cluster. Similar trends were observed in regions containing *RPP2A/B, WRR4*, and especially *RPP7* (*SI Appendix*, Table S10-12 and Fig. S5), showing that CES-mediated methylation changes are part of a broader, more global effect that is not restricted to the *RPP5* gene cluster.

### CES Associates with Chromatin Remodelling and Splicing Factors

CES is a bHLH TF which in response to BR signaling activation is SUMOylated, yielding an accumulation of the protein in subnuclear compartments ^21^. CES SUMOylation at K72 is inhibited by phosphorylation at two serines (S75 and S77) in close vicinity to the SUMO site; accordingly, a CES mutant in which both serines are exchanged to alanines (CES^S75A+S77A^; CES_AA_) is constitutively enriched in subnuclear domains ^21^.

The nature of CES-containing nuclear bodies has remained unknown and we speculated that they may contain proteins that facilitate CES-mediated DNA methylation changes. To test this idea, YFP-tagged version of CES_AA_ and CES_wt_ were used as bait for a co-immunoprecipitation (co-IP) of associated proteins and their identification with liquid chromatography coupled to tandem mass spectrometry (LC-MS/MS). A total of 944 CES-associated proteins were pulled (*SI Appendix*, Table S13) and classified based on their annotation to cellular components with the GO Ontology (GO) tool. The most significantly enriched GO groups entailed the terms ‘mRNA binding’, ‘RNA binding’, ‘structural molecule activity’ and ‘structural constituent of ribosomes’, among the top 20 categories (Fig. 3*A*; *SI Appendix*, Table S14). Within these most enriched categories, multiple subunits of the SWI2/SNF2 (SWR1) chromatin remodeling complex were present, namely SWR1 COMPLEX 4 (SWC4), ACTIN-RELATED PROTEIN 4 (ARP4), RUVBL1 (RIN1), and RUVBL2A (RIN2), which regulates chromatin structure through histone H2A.Z deposition ^38,39^ and promotes active DNA demethylation through the recruitment of the DNA glycosylase ROS1 ^40^.

**Fig. 3.**
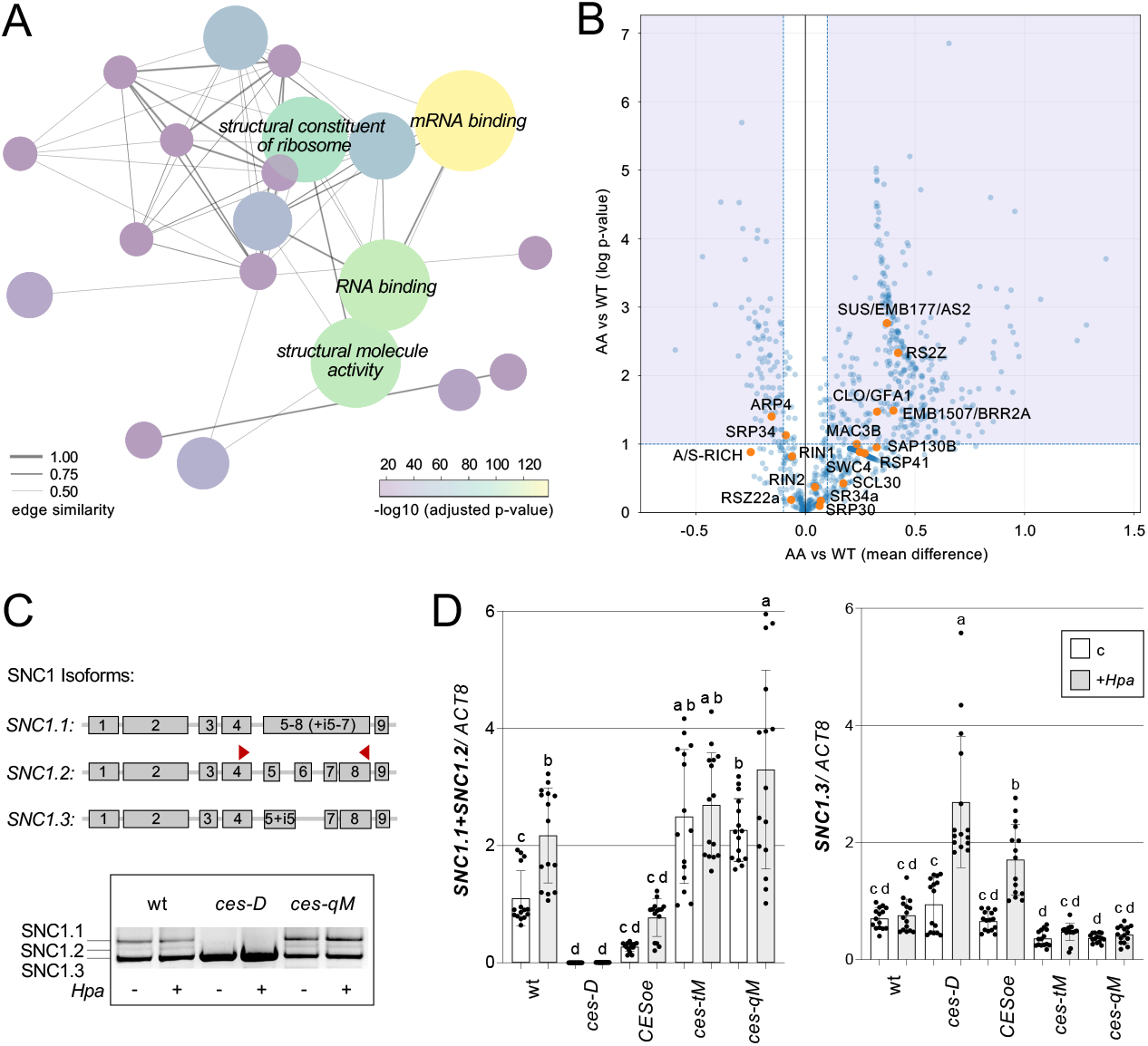
CES co-localizes with chromatin remodeling and splicing factors and modulates *SNC1* mRNA splicing. (*A*) Molecular function enrichment map of proteins co-purifying with CES-YFP, identified by GCMS/MS. Highly enriched categories include components of the SWR1 chromatin remodeling complex, the MAC, and the spliceosome. Representative components are listed in the accompanying table. (*B*) Vulcano blot comparing the abundance of proteins pulled with either CES_AA_-YFP or CES_wt_-YFP as baits. Components of the SWR4 chromatin remodeling complex, MAC and spliceosome are highlighted in orange. (*C*) Top: Scheme of predominant *SNC1* mRNA isoforms in wt and *ces* mutants. Primer positions used to distinguish the three isoforms are indicated by red arrows. Bottom: Detection of alternatively spliced *SNC1* transcripts by semiquantitative RT–PCR in wt, *ces-D*, and *ces-qM* plants, either untreated (-) or infected with *Hpa* (+), using the primers shown in *B*. (*D*) Absolute expression levels of *SNC1* isoforms (.1, .2, and .3) measured by qPCR in *ces* mutants and wt, untreated or infected with *Hpa*. Data represent mean ± SD from four biological replicates, each measured in triplicate and normalized to *ACT8*. Statistical significance was assessed using one-way ANOVA with Tukey’s HSD post hoc test in GraphPad Prism v10. All tests were two-sided; *P* < 0.05 was considered significant.

In addition, and very interestingly, a large set of proteins involved in pre-mRNA splicing were identified, such as the SPLICEOSOME-ASSOCIATED PROTEIN 130B (SAP130B) and the serine/arginine-rich splicing factors SR30, SR34, SR34a, RSZ22a, RS2Z32, RS2Z33, RS41, and SCL30. Moreover, several components of the MOS4-associated complex (MAC) were enriched, namely MAC3B, CLOTHO/ MATERNAL EFFECT EMBRYO ARREST 5 (CLO/ MEE5), EMBRYO DEFECTVE 1507/ BRR2 HOMOLOGUE 2 (EMB1507/ BR22A) and ABNORMAL SUSPENSOR 2 (SUS2/ EMB177). The MAC is an evolutionarily conserved complex that acts in the catalytic activation of the spliceosome and is well-known for its function in *SNC1* pre-mRNA splicing ^41,42^.

Given that splicing factors, including MAC components, are concentrated in nuclear bodies ^43,44^, BR-induced CES nuclear body enrichment likely promotes its association with them. In line with this idea, most pulled MAC components were significantly more enriched when the constitutively speckling CES_AA_ mutant was used as a bait for co-IPs (Fig. 3*B; SI Appendix*, Table S13).

Chromatin Immunoprecipitation (ChIP) experiments showed that CES actively bound to the *SNC1* locus region (*SI Appendix*, Fig. S6), and it seemed possible that it may associate with the spliceosome to alter *SNC1* splicing. To test this, the abundance of *SNC1* splice variants in wt and *ces* mutants was quantified. Sequencing of *SNC1* cDNAs (*SI Appendix*, Fig. S7) showed that three major types with variations 3’ of exon 4 were formed in the *ces* mutants and their wt Col-0: a strongly spliced form (*SNC1*.*2*), a transcript that retained introns 5, 6 and 7 (*SNC1*.*1*; the main isoform reported in TAIR), and a variant with partial retention of intron 5 and skipping of exon 6 (*SNC1*.*3*) (Fig. 3*C*). Notably, the levels of *SNC1*.*1* and *SNC1*.*2* were significantly altered in the *ces* mutants compared to wt, with *ces-D* exhibiting a strong decrease and *ces-qM* an increase (Fig. 3*C, D*). These changes shifted the splice variant distribution, resulting in a predominance of *SNC1*.*3* in *ces-D* and of *SNC1*.*1+SNC1*.*2* in *ces-qM*, an effect that was further amplified by *Hpa* treatment. The *CESoe* and *ces-tM* lines showed similar trends but more subtle outcomes, consistent with the results from our other analysis. Importantly, the over-all mRNA abundance of *SNC1* was not consistently altered in all *ces* mutants (*SI Appendix*, Fig. S8), demonstrating that CES regulates *SNC1* predominantly at the level of alternative splicing, thereby explaining why RNA-seq did not detect significant changes in *SNC1* expression. In contrast, the previously published microarray analysis ^20^ likely employed probes specific for the *SNC1*.*3* isoform, thereby capturing isoform-specific changes.

To investigate if CES also regulates other NLRs, two additional members of the RPP5 cluster where DNA methylation changes in *ces* mutants occurred, *RPP4*, and *At4g16900*, were tested. This analysis revealed altered splicing outcomes in CES *gof* and *lof* backgrounds for both genes, but more strongly for *At4g16900* (*SI Appendix*, Fig. S9*A, B*), supporting the conclusion that CES-dependent regulation of alternative splicing is not limited to *SNC1*.

### BRI1 Acts Up-stream of CES in *SNC1* Splicing

CES enrichment in subnuclear compartments is promoted by BRs ^21^ and it was thus relevant to investigate if BR signaling is required for CES-induced alternative splicing. Therefore, the *ces-D* effects on *SNC1* splicing were assessed in the *bri1-1* null mutant background, employing a *ces-D×bri1-1* double mutant ^24^. This showed that the decreased abundance of *SNC1*.*1+2* in *ces-D* was partially restored in *bri1-1* providing evidence that CES function in *SNC1* splicing is promoted by BR perception (*SI Appendix*, Fig. S10*A*).

To explore if BRI1 also impacts DNA methylation, WGBS was performed with *bri1-1* and the triple mutant *bri1 brl1 brl3 (brl-tM)*, in which expression of BRI1 and its two homologs BRI1-like 1 and 3 is lost ^45^. The DMR profiles showed that the *BRI1* mutation induced strong global changes in DNA methylation, which were highly similar to those of the *ces* mutants (*SI Appendix*, Fig. S10*B*). A comparison of hyper- and hypo-methylated DNA regions was performed and showed that hypo-methylated area shared significant portions (χ^2^ test, *p*=2.62e-47) and that these were common not only between both BR-receptor mutants, but also with *ces-qM* (*SI Appendix*, Fig. S10*C*).

Significance was given in pulling the overlap of DMRs shared in the hypo CHH context (χ^2^ test, *p*=0.02) but more so for the hyper CHH contexts (χ^2^ test, *p*=1.92e-66) (*SI Appendix*, Fig. S10*D*).

Although the global methylation profiles of *bri1-1* and *brl-tM* mutants closely resembled those of *ces-qM*, the specific NLR loci analyzed as representative readouts, namely those containing *RPP5* (*SI Appendix*, Fig. 10*E*) and *RPP7* (*SI Appendix*, Fig. S11), exhibited more subtle changes compared to *ces-qM*. While all three mutants shared DMRs at similar coordinates, the changes in the *bri1* mutants were milder and more confined to CHH methylation changes in TE-containing areas.

To directly assess whether CES is required for BR-induced DNA methylation reprogramming, BL treatments with wt, *ces-D* and *ces-qM* were performed, followed by WGBS of untreated and treated samples. In wt, BL-treatment induced widespread changes in DNA methylation across the genome (*SI Appendix*, Fig. S12). Importantly, this BR-responsive methylation landscape was markedly altered in both *ces-D* and *ces-qM*. The differences were most apparent in the abundance and genomic distribution of hyper- and hypo-methylated regions, particularly in CG and CHH contexts, supporting the conclusion that BR-induced methylome reprogramming requires CES.

Changes in the RPP5 locus-specific methylation profile following BL treatment also mirrored distinct differences as compared to wt in both *ces-D* and *ces-qM*. In the *gof ces-D* mutant, methylation patterns were characterized by broad and persistent hypo-methylation across TE-containing regions. By contrast, in the *ces-qM lof* background, where more discrete changes would be expected, BL-treatment resulted in fewer, well-defined DMRs, predominantly in the CG and CHH context (*SI Appendix*, Fig. S13, Table S15). In both mutants, CHH methylation was consistently altered at the same NLR-proximal TEs, showing that CES is required to properly establish BR-induced methylation changes within the RPP5 cluster.

### ces-D Represses Autoimmunity in snc1

Since CES had largescale repressive effects on immunity-related genes, it was relevant to assess if this activity can be utilized to limit growth suppression. To test this, *ces-D* was introduced into *snc1* by crossing, resulting in a *ces-D×snc1* double mutant in which *CES* transcript levels were comparably elevated like in the *ces-D* parent (Fig. 4*A*). *snc1* contains a gain-of-function, single point mutation that leads to over-accumulation and constitutive activation of SNC1 ^28^. An increased CES activity in *ces-D×snc1* resulted in the suppression of the generation of *SNC1*.*1*+*SNC1*.*2* isoforms (Fig. 4*A*), and a full recovery of the constitutive *Hpa* resistance (Fig. 4*B*) and dwarfism of *snc1* (Fig. 4*C*). This shows that isoforms *SNC1*.*1+SNC1*.*2* are essential for conferring *Hpa* resistance and growth repression in *snc1* and that CES can reduce their abundance, thereby limiting fitness cost. In support of this idea, it happens that CES-induced splicing truncates the LRR domain of SNC1 (*SI Appendix*, Fig. S14), which in NLRs is required for effector recognition ^46^.

**Fig. 4.**
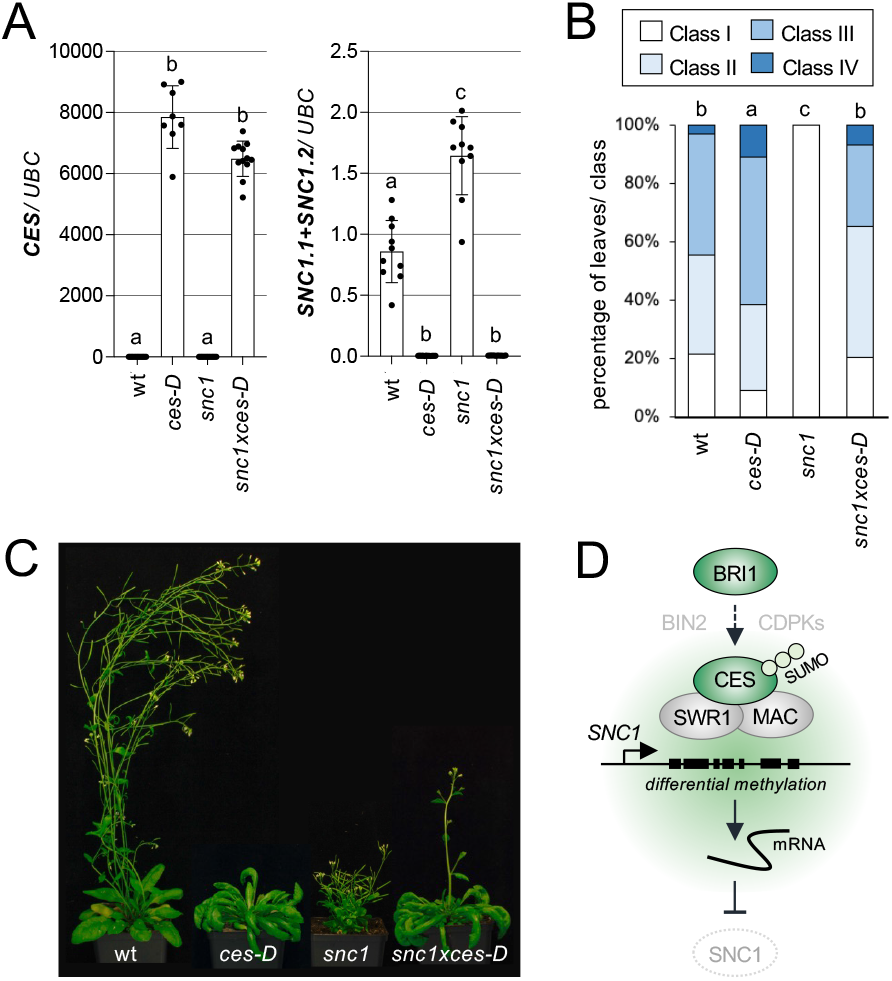
*ces-D* represses *SNC1*.*1* and *SNC1*.*2* isoform formation and autoimmunity in *snc1*. (*A*) Absolute expression levels of *CES* (left) and *SNC1*.*1*+*SNC1*.*2* (right) in 18-day-old plants of the *snc1×ces-D* cross and parental lines, measured by qPCR. Data represent mean ±SD from four biological replicates, each measured in triplicate and normalized to *UBC*. (*B*) *Hpa* colonization in the same genotypes as in a. Two-week-old plants were spray-inoculated with *Hpa* Noco2, and colonization was assessed microscopically at 6 dpi in trypan blue-stained leaves. Infection was categorized into four classes; data represent the percentage of leaves per class (n = approx. 100 leaves from 20-25 plants per genotype). (*C*) Representative photos of five-week-old plants of the *snc1×ces-D* cross and parental lines grown under long-day conditions. (*D*) Proposed model of CES function in chromatin remodeling and alternative splicing of *SNC1*. CES initiates DNA methylation changes and alternative splicing of SNC1 to repress its activity. This process entails association with chromatin remodeling and splicing factors including the MAC, and CES enrichment in subnuclear compartments.

## Discussion

The steroid hormones BRs are known repressors of plant immunity, with effects on resistance against a range of pathogens including *Hpa* ^3^. Here we expand our understanding of their modes of activity in this process, by revealing that the BR-controlled TF CES effects DNA methylation at a global level and alters splicing of the NLR *SNC1*.

We show that BRs, via CES, induce DNA methylation reprogramming, which is characterized by CHH hypo-methylation in TE-rich regions and CG hyper-methylation at adjacent genes. These changes prominently affect the *RPP5* gene cluster, including its member *SNC1*. CES binds chromatin upstream of *SNC1* and associates with the SWR1 chromatin remodeling complex, which is known to recruit the DNA glycosylase ROS1 to demethylate specific loci ^40^. This suggests CES may facilitate active demethylation through SWR1-ROS1 recruitment.

Alternatively, SWR1-mediated H2A.Z deposition may itself antagonize DNA methylation via histone acetylation ^46^. The process may be promoted by SUMOylation, which not only targets CES for an enrichment in nuclear bodies, but also promotes ROS1 activity and active DNA demethylation in *A. thaliana* ^23^.

CES-mediated DNA methylation changes extended from intergenic regions into gene bodies, and there is evidence that methylation of gene bodies can influence splicing, for example by preventing aberrant intron retention ^9^. Moreover, results from yeast showed that H2A.Z. deposition through SWR1 is linked to pre-mRNA splicing ^47,48^ and here we show that CES-mediated methylation changes upstream and within the *SNC1* gene correlated with alternative splicing of this immune receptor. In a *ces* knock-out situation, *SNC1*.*1* and *SNC1*.*2* isoforms were over-accumulated, whereas *CES* over-expression suppressed their formation. In addition, CES associates with spliceosome components including the MAC, previously implicated in *SNC1* regulation ^42^, suggesting CES directly modulates pre-mRNA processing, creating shifts in splice variant ratios, a process that is known to effect the activity of NLRs, including SNC1 ^50,51^. Given that the MAC promotes SNC1 activity ^40^, CES would be expected to impair MAC function and it will be important to genetically test this idea. Altered splicing of *SNC1* and additional CES targets such as *RPP4* and *At4g16900* may also feed into DNA methylation changes in the *ces* mutants, since genotypes with splicing defects, and in particular MAC component mutants, display defective RdDM ^52^.

These findings suggest a model where CES acts as a key integrator of BR signaling, chromatin state, and resistance gene expression (Fig. 4*D*). In response to BRs, CES initiates DNA methylation changes and alternative splicing of immunity-related genes including, but not restricted to, NLRs such as *SNC1*, a process that entails SUMOylation-induced nuclear body enrichment of CES. This could restrain NLR activity under stress-free conditions, but also facilitate immune resetting post-activation, thereby minimizing autoimmunity and associated energy costs. Notably, this regulation may also occur by BRI1-independent means, for example through spatial control of CES post-translational modifications and localization. For instance, CES phosphorylation at S75+S77 by CDPK3, a calcium-dependent protein kinase activated upon pathogen recognition that contributes to both PTI and ETI ^53^, prevents SUMOylation and CES nuclear body enrichment ^21^, which may link immune signaling to chromatin remodeling and alternative splicing. In addition, CES transcription is known to be repressed by PAMPs ^17^, and it will be interesting to further investigate this potential multilayered control of CES activity by immune signaling.

Because CES loss-of-function does not trigger constitutive immune responses but rather finetunes and sensitizes them, it appears to function in priming, adjusting epigenetic and splicing states to promote immune responses while avoiding growth penalties. Harnessing this pathway, by tuning CES activity or localization, could enable breeding of crops with enhanced resistance and minimized growth-immunity trade-offs.

## Materials and Methods

### Plant Material and Growth Conditions

All single mutants and over-expression lines used in this study were in the Col-0 background and have been previously described ^17,20,25,28,45^. The double mutant *ces-Dxsnc1* was generated by crossing the respective single mutants and selecting homozygous progeny through genotyping (all primer sequences are provided in SI Table 14). For phenotypic analyses and infection assays, seeds were germinated on ½ MS medium (Duchefa Biochemie, Haarlem, The Netherlands), transferred to soil 7-8 days after germination and grown under long-day conditions (16 h light/8 h dark, 80 μmol m ^2^ s ^1^, 50% humidity) in a controlled environment growth cabinet (Bright Boy, CLF Plant Climatics, Wertingen, Germany) maintained at 21°C +/-2 for the required amounts of time.

### *Hpa* Infection Assays and Resistance Scoring

*Hpa* infection assays were performed as described previously ^54^. In brief, the *Hpa* isolate Noco2 was propagated on wt *A. thaliana* Col-0 plants, leaves were harvested, placed in tubes with double-distilled water and vortexed. The suspension was then filtering through Miracloth (Millipore-Sigma, Burlington, MA, USA), and the spore concentration was adjusted to 5 × 104 conidiospores/ mL using a hemacytometer (Paul Marienfeld GmbH, Lauda-Königshofen, Germany). Two-week-old plants were inoculated by spraying with the spore suspension or water as a control using a standard atomizer. Following inoculation, plants were placed in sealed trays and incubated at 18°C under 16 h light/8 h dark photoperiod and a relative humidity of 90–100%.

For resistance scoring, infection was assessed at 5 dpi by counting spores with a Fuchs Rosenthal counting chamber and at 7 dpi by assessing sporangiophore formation following trypan staining. For this purpose, infected aerial tissue (≥30 plants) was stained with Trypan Blue (Sigma-Aldrich, St. Louis, MO, USA) by heating at 95°C for 1 min and incubating for 1 h at room temperature. Samples were cleared overnight in chloral hydrate (Sigma-Aldrich), then incubated in 60% glycerol (Sigma-Aldrich). Stained tissue was rinsed and stored in 80% glycerol prior to quantification. Sporangiophores were counted using an Olympus SZX12 stereomicroscope (Olympus, Tokyo, Japan). Resistance phenotypes were classified according to ^10^.

### ROS Burst Assays

ROS burst assays were performed with leaf discs from 4-5-week-old plants using 100 nM flg22 or elf18 as described previously ^55^ and luminescence was quantified using a charge-coupled device camera (Photek Ltd, East Sussex, United Kingdom).

### SA Measurements

For SA measurements, approximately 200 mg of 18-day-old seedlings were flash-frozen together with 2.8 mm zirconium beads (BioSpec Products, Bartlesville, OK, USA) and homogenized. Each sample was spiked with 20 μL of isotope-labeled SA (Merck, Darmstadt, Germany) in acetonitrile as an internal standard. Metabolites were extracted using glacial ethyl acetate (Sigma-Aldrich) with a Percellys Homogenizer (Bertin Technologies, Montigny-leBretonneux, France). The resulting supernatant was filtered through a 0.45 μm membrane (Sartorius, Göttingen, Germany) and evaporated to dryness. Dried extracts were resuspended in 70 μL acetonitrile (Sigma-Aldrich) and 2 μL were injected for LC-MS/MS, performed as described previously ^56^. 3-4 biological repeats per genotype were analyzed and the results were evaluated by two-sided unpaired Student’s t-tests.

### Semi-Quantitative PCR and Real-Time qPCR Analyses

For gene expression analyses by semi-quantitative PCRs and real-time qPCRs, plants were grown and infected as described above and harvested at 7 dpi. Total RNA was extracted from 100 mg of tissue using the E.Z.N.A. Plant RNA Kit (Omega Bio-Tek, Norcross, GA, USA) and quantified with a NanoDrop 2000 spectrophotometer (Thermo Fisher Scientific, Waltham, MA, USA). Two micrograms of total RNA were treated with DNase I (Thermo Fisher Scientific) and reverse transcribed using a combination of oligo(dT) and random hexamer primers, dNTPs (Thermo Fisher Scientific), and M-MuLV Reverse Transcriptase (Thermo Fisher Scientific), following manufacturer’s instructions. Semi-quantitative PCRs were performed by PCR by amplifying the isoform-resolved C-terminal sequences of *SNC1, RPP4, and At4g16900* from 2 μL of cDNA with primers listed (*SI Appendix*, Table S16) and Phusion High-Fidelity DNA Polymerase (Thermo Fisher Scientific) and separating the products on a 1.5% (w/v) agarose gel. Band identity and intensity was used for qualitative comparison of *SNC1* isoform abundance between lines and treatment conditions.

RT-qPCRs were performed in 10 μL reaction volumes using qPCRBIO SyGreen Mix (PCR Biosystems, London, UK) on a Bio-Rad CFX96 cycler (Bio-Rad, Hercules, CA, USA). Each reaction contained 5 μL of 2x SyGreen mix (Nippon Genetics), 0.4 μL each of forward and reverse primers (10 μM), 4 μL cDNA, and 0.2 μL nuclease-free water. Gene expression levels were normalized to the reference genes *UBIQUITIN* (*UBC*) ^57^, *ACTIN8* (*ACT8*) ^58^, or *TUBULIN* 2 (*TUB2*). The analyses were performed with at least three biological replicates per genotype and treatment condition each measured in technical triplicates, and all replicates were used for data analysis. Statistical significance was assessed in GraphPad Prism v10 using one-way ANOVA to compare expression levels across multiple genotypes and treatments, followed by post hoc multiple comparison testing (e.g., Tukey’s HSD or Bonferroni) where appropriate. All tests were two-sided; significance P < 0.05.

### ChIP Experiments

For ChIP experiments, plants of *35S:CES-YFP* line 32 ^20^ were grown on ½ MS medium under long-day conditions for three weeks. They were then sprayed with 10 μM 24epibrassinolide (Sigma-Aldrich) and four biological replicates each containing approximately 400 seedlings, or two grams of tissue, were harvested 3 hours post-treatment. Chromatin crosslinking and extraction were performed as previously described ^24^. Immunoprecipitation was carried out using magnetic anti-GFP VHH agarose beads (ChromoTek, Planegg-Martinsried, Germany) with control reactions performed using non-specific agarose beads (ChromoTek). DNA was purified with MinElute Reaction cleanup kit (Qiagen, Hilden, Germany). Promoter fragments of *SNC1* were quantified by qRT-PCR, and values were normalized to blocked bead background and calculated as input percent ratios compared to wt. ChIP was performed in three biological replicates with four technical repeats each. Enrichment was calculated as the ratio of antibody treated to untreated signal. Statistical comparisons were made using two-sided Student’s t-tests.

### Sequencing of Splice Variants

2 μL of cDNA from wt and *ces-D* was used as template in PCR reactions with Phusion High-Fidelity (Thermo Fisher Scientific, Waltham, MA, USA) and the respective primers (*SI Appendix*, Table S16) were used for sequencing. The amplicons were separated on a gel, the different isoform fragments were extracted, sequenced, and the sequences were aligned against the respective reference TAIR10 CDS sequence using SnapGene Viewer (Dotmatics). Alignments were verified by visual inspection and refined using Clustal Omega (EMBL-EBI).

*RNA-seq, WGBS, mass spectroscopy-based proteomics and Co-IP, as well as statistical analyses, see SI Appendix*.

## Supporting information

Ramirez et al., 2026 Supporting Information

Ramirez et al., 2026 Supplementary Tables

## Acknowledgments

We thank Jian Hua (Cornell University) for kindly providing seeds of *snc1*. This work was supported by funding from the Elite Network of Bavaria (grant F-6-M5613.6.K-NW-2021-411/1/1 to B.P., C.D., C.G., B.K. and C.L.), the German Research Council DFG (Collaborative Research Center SFB924 project B03 to C.G.), the University of Zürich (C.Z.) and the Swiss National Science Foundation (grant no. 310030_212382 to C.Z.), and the China Scholarship Council (Ph.D. fellowship to H.S.). The Orbitrap Fusion Lumos Tribrid mass spectrometer was funded in part by the German Research Foundation (INST 95/1436-1 FUGG).

