## Supplementary material for "Brassinosteroid-regulated transcription factors confer epigenetic changes that repress plant immunity": Ramirez et al., 2026 Supporting Information

**This PDF file includes:**

**Supplementary Materials and Methods**

**Supplementary Figures**

**Figure S1.** SA levels in *ces* mutants are not indicative of autoimmunity.
**Figure S2.** RNA-Seq data analysis.
**Figure S3.** Overexpression of *HBI1* neither significantly alters *SNC1* splicing nor *Hpa* Noco2 resistance.
**Figure S4.** CES represses ROS responses.
**Figure S5.** CES confer DNA methylation changes in different NLR-containing chromosome region.
**Figure S6.** CES can bind to the *SNC1* promoter *in vivo*.
**Figure S7.** The *SNC1* mRNA is alternatively spliced.
**Figure S8.** Overall *SNC1* transcript abundance in *ces* mutants.
**Figure S9.** CES modulates *RPP4* and *AT4G16900* splicing.
**Figure S10.** *BRI1* loss-of-function confer *ces* mutant-like patterns of DNA methylation changes.
**Figure S11.** *bri1* mutants show DMRs in TE-containing regions of the *RPP7* locus region.
**Figure S12.** BRs induce global DNA methylation changes that are altered in *ces* mutants.
**Figure S13.** CES mutation alters BR-induced DNA methylation changes.
**Figure S14.** AlphaFold-based structural modeling of SNC1 isoforms.

**Other supporting materials for this manuscript include the following:**

Tables S1-S16

### Materials and Methods

**RNA-seq Analysis.** For RNA-seq, RNA was extracted from 3-week-old mock-treated or *Hpa*infected Arabidopsis plants in five biological replicates using the E.Z.N.A. Plant RNA Kit (Omega Bio-Tek, Norcross, GA, USA) with the difficult sample protocol in the manufacturer's instructions.
RNA RIN quality was assessed using the 2100 Bioanalyzer (Agilent Technologies, Santa Clara, CA, USA). Libraries were prepared using a modified SCRB-seq protocol [1] similar to prime-seq [2] with barcoded oligo-dT primers (IDT, Coralville, IA, USA) and reverse transcription using KAPA HiFi HotStart polymerase (Roche/KAPA Biosystems, Wilmington, MA, USA). Tagmentation and library PCR were performed with Nextera XT DNA Library Prep Kit (Illumina, San Diego, CA, USA). Libraries were size-selected on 2% E-Gel EX Agarose Gels (Thermo Fisher Scientific). Sequencing was done on an Illumina HiSeq1500 (Illumina), and data were processed using zUMIs [3].

Analysis was performed using DESeq2, PANTHER (2024 release), and custom Python scripts utilizing Matplotlib, Plotly, NumPy, SciPy, and Pandas. DEGs were identified using adjusted $P < 0.05$ and $\log_2 FC \geq 1$. For visualization, normal expression values (VST from DESeq2) were used tom compare expression changes between *Hpa*-treated and mock-treated samples within genotypes. Hierarchical clustering was performed on these values to group genes and samples based on expression similarities, revealing distinct patterns within the lines following *Hpa* treatment.

Gene Ontology (GO) term enrichment was performed using the PANTHER database and Fisher's exact test with FDR correction (adjusted $P < 0.05$). Raw and processed RNA-seq data have been deposited in the NCBI Gene Expression Omnibus (GEO) repository under accession number GSE304560. Access is available upon request/provided to editors and reviewers.

**WGBS Analysis.** DNA from five biological replicates of 100 mg aerial tissue of 3-week-old, soil grown plants was extracted using the DNeasy Plant Mini Kit (Qiagen, Hilden, Germany). For eBL treatments, either 10µM 2,4-epi-brassinolide or DMSO (control) was sprayed exogenously on seedlings, and material snap frozen in LN after 3h. DNA precipitation was performed with 3 M sodium acetate (Sigma-Aldrich), 100% ethanol (Sigma-Aldrich), and glycogen (Thermo Fisher Scientific). Samples were either sent to BGI Genomics (Tech Solution NGS Lab, Hong Kong, China) for bisulfite conversion and sequencing using DNBseq on the Illumina X Ten platform, or to Novogene, (Novogene GmbH, Munich, Germany) for library prep and sequencing on the Illumina NovaSeq X Plus Series (PE150). Raw WGBS reads were processed with the MethylStar pipeline (v1.2.3). Initial quality assessment was performed using FastQC, and adapter and lowquality base trimming was carried out with Trim Galore. Cleaned reads were aligned to the *A. thaliana* TAIR10 (v57) reference genome using Bismark (v0.23.0) in combination with Bowtie2. Cytosine methylation states were called with METHimpute (v1.18.0) [4] providing probabilistic estimates for CG, CHG, and CHH sequence contexts. Strand-specific methylation calls were then converted into bigWig format using custom in-house scripts.

Context-specific DMR analysis of the *ces-D*, *ces-qM*, *bri1-1*, and *brl-tM* mutants relative to wt was performed using jDMR [5], which segments the genome into non-overlapping 100 bp bins. All bins were included in the analysis, and DMRs were defined using the built-in control/treatment mode, requiring replicate consensus (set to 1) to filter for reproducible differences. To avoid batch effects, each mutant was compared to its designated control wt group, with separate wt controls used for the *ces-D/ces-qM* and *bri1-1/brl-tM* mutant groups. All genomic coordinates refer to the TAIR10 genome build. For genome-wide comparisons, DMR counts were computed per context and genotype, and distribution profiles were visualized without prior smoothing. In addition to the *ces* and *bri1* mutants, public datasets from PRJNA172021, PRJNA176484, and PRJNA842430, containing *ago4-5*, *cmt3-11*, *drm2-3*, *nrpe1-11*, and *ros1-4* methylation values [6-8], were analyzed with this approach.

Strand-specific methylation bubble plots were generated from bigWig-formatted cytosine methylation data using custom Python scripts (pyBigWig v0.3.18, Pandas, Matplotlib v3.7) in a Jupyter Notebook environment. Methylation signals for each context were extracted from defined genomic intervals, filtered to remove negative values, and plotted as scatter plots with bubble size indicating methylation density. Replicates were visualized side-by-side by genotype, with consistent DMRs across biological duplicates retained for further analysis. No additional statistical testing was applied to plots; METHIMPUTE-derived genome-wide DMR analyses supported reproducibility.

WGBS data have been deposited in the NCBI GEO repository under project numbers GSE304334, GSE325577, and GSE325577. Access is available upon request/provided to editors and reviewers.

**Mass Spectrometry-Based Proteomics.** For proteomics aerial plant tissue from *Hpa*-treated and mock-treated plants was lysed via TCA/acetone (Sigma-Aldrich) precipitation, washed with acetone (Sigma-Aldrich), and dissolved in SDS buffer. Proteins were extracted with TE-buffered phenol (Sigma-Aldrich), precipitated with ammonium acetate in methanol (Sigma-Aldrich), and quantified by BCA assay (Thermo Fisher Scientific). SP3 beads (Sera-Mag SpeedBeads; Cytiva, Marlborough, MA, USA) were used for protein cleanup and digestion. Peptides were desalted with Chromabond HLB plates (Macherey-Nagel, Düren, Germany). MS analysis was performed on an Orbitrap Fusion Lumos Tribrid with a Vanquish Neo UHPLC system and PepMap C18 column (both Thermo Fisher Scientific).

Data were analyzed with MaxQuant v2.1.0.0 and Perseus with FDR = 0.01 and $s_0 = 0.5$. Analyses and resulting heatmaps were generated $log_2$-transformed values using Python (v3.9) with the seaborn (v0.11.2), pandas (v1.5.3), and matplotlib (v3.5.1) libraries. Raw abundance values were Z-score normalized across samples and clipped (–3 to +3). Heatmaps were plotted using seaborn.clustermap, with hierarchical clustering enabled for rows and disabled for columns. Clustering was performed using the Euclidean distance metric and average linkage. Chi-squared tests were used to assess overlap between protein and transcriptomic datasets ($P < 0.05$), with analyses performed in Python.

Raw and processed proteomics data have been deposited to the ProteomeXchange Consortium via the PRIDE partner repository and can be accesses with the identifier PXD066892. Access is available upon request/provided to editors and reviewers.

**Co-Immunoprecipitation and Identification of CES-Associated Factors.** Seeds of the lines 35S:CES$^{wt}$-YFP/32 and 35S:CES$^{S75A+S77A}$-YFP/310 [9,10] were germinated on ½ MS medium and after 10 days approximately 2 g of plant tissue per line were harvested and ground to a fine powder in liquid nitrogen. The powder was resuspended in 6 mL of lysis buffer (50 mM Tris/HCl pH 7.5, 150 mM NaCl, 10 mM $MgCl_2$, 10% glycerol, 1 mM EDTA, 5 mM DTT, 0.2% Nonidet$^{TM}$ P40, 1 mM PMSF, 1x protease inhibitor cocktail), vortexed for 1 min and incubated on ice for 30 min. Cell lysates were centrifuged at 2,000 × g for 15 min at 4 °C, and the cleared supernatant was collected. Total protein concentration was determined using the Bradford Assay.

50 µL GFP-Trap agarose beads (ChromoTek) were prewashed with 500 µL of lysis buffer and incubated with supernatant containing 5 mg of total protein at 4 °C for 2 h with gentle rotation. Beads were then pelleted by centrifugation at 2,500 × g for 5 min at 4 °C, and the supernatant was discarded. The beads were washed twice with 500 µL of lysis buffer, were resuspended in 80 µL of 2x SDS-sample buffer (120 mM Tris/HCl pH 6.8, 20% glycerol, 4 % SDS, 0.04% bromophenol blue, 10% 2-mercaptoethanol) and boiled at 95 °C for 5 min. The supernatant was collected after centrifugation and an in-gel trypsin digestion and mass spectrometric measurements were performed as described previously [11] with the LC-MS/MS data acquisition carried out on a Dionex Ultimate 3000 RSLCnano system coupled to an Orbitrap Fusion Lumos Tribrid mass spectrometer (Thermo Fisher Scientific).

Peptide identification and quantification was performed using the software MaxQuant (version 1.6.3.4) [12]. MS2 spectra were searched against the *A. thaliana* protein database from TAIR, supplemented with common contaminants (built-in option in MaxQuant). Results were adjusted to a 1% FDR on peptide spectrum match and protein level employing a target-decoy approach using reversed protein sequences. Label-Free Quantification [13] was used for protein quantification with at least 2 peptides per protein. The minimal peptide length was defined as 7 amino acids and the "match-between-runs" functionality was disabled.

To explore the functional enrichment of gene sets, GO analysis was performed using g:Profiler(https://biit.cs.ut.ee/gprofiler. Differentially expressed proteins obtained from MQCruncher were submitted to the g:GOSt module, specifying *A. thaliana* (TAIR/Ensembl Plants) as the organism. Analyses were conducted against the GO: Molecular Function (MF) namespace, using default g:SCS multiple testing correction to control for FDR. Enrichment results were exported in matrix format, containing term IDs, names, adjusted p-values, gene intersections, and associated statistics. Only categories with adjusted $P < 0.05$ (Benjamini–Hochberg correction) were retained. The resulting dataset was processed and visualized in a custom pipeline implemented in Python 3.11 within a Jupyter Notebook environment. The GO terms were filtered to retain only the top 20 most significantly enriched terms, based on adjusted p-values. For each selected GO term, the list of intersecting genes was extracted from the "intersections" column. Pairwise functional similarity between terms was estimated using Jaccard similarity, calculated as the proportion of shared genes over the union of genes between any two terms.

An undirected network was constructed using the NetworkX library (v2.8), where nodes represent GO terms and edges denote functional similarity (Jaccard index ≥ 0.5). Node size was scaled according to the term significance (−log10 adjusted p-value), and node color was mapped to the same metric using a pastel-modified version of the *viridis* colormap for improved legibility and visual appeal. Edges were drawn with line width proportional to similarity (range: 0.5–4.0 points), with thicker edges indicating greater gene overlap. All visual elements were rendered using matplotlib (v3.7), with layout positions determined via a force-directed spring layout (spring_layout, seed=42).

The mass spectrometric raw files as well as the MaxQuant output files have been deposited to the ProteomeXchange Consortium via the PRIDE partner repository: identifier PXD067053. Access is available upon request/provided to editors and reviewers.

**Statistics and Reproducibility.** Statistics were derived from data of biological replicates with technical repeats. Pairwise comparisons of spore/sporangiophore counts among mutant genotypes performed using two-sided Mann–Whitney U-tests. Missing data points were excluded on a pairwise basis. Analyses were either conducted with GraphPad Prism v10 and confirmed using Python's SciPy.Stats package. Significant differences were identified based on $P < 0.05$. Where applicable, p-values were corrected for multiple comparisons using the Bonferroni method. Trypan Blue staining results were categorized into four qualitative classes based on sporulation and tissue response. To assess the statistical significance of differences in phenotypic class distributions across genotypes, pairwise Fisher's exact tests were performed using a 4×2 contingency table for each genotype pair. Multiple comparisons were corrected using the Benjamini–Hochberg FDR method, and adjusted P-values are reported. Statistical significance was defined as $p < 0.05$ after correction. Significantly distinct genotype groupings were labeled using compact letter display to indicate statistically non-overlapping classes. Analyses were conducted using standard statistical packages in R (e.g., fisher.test and p.adjust) or Python (scipy.stats.fisher_exact with FDR correction via statsmodels) unless otherwise indicated.

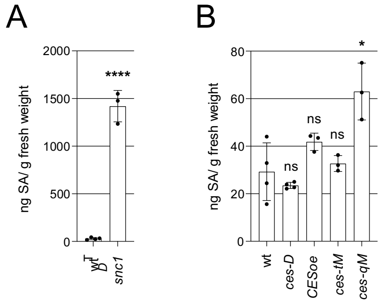


**Fig. S1. SA levels in *ces* mutants are not indicative of auto-immunity.**

SA levels (ng/ g fresh weight) in 18-day-old plants of *snc1* (*A*) and *ces* mutants (*B*) as compared to wt, quantified by UHPLC-MS/MS. Statistical significance was determined with Students T-test (ns P > 0.05, *P ≤ 0.05, ****P ≤ 0.0001).

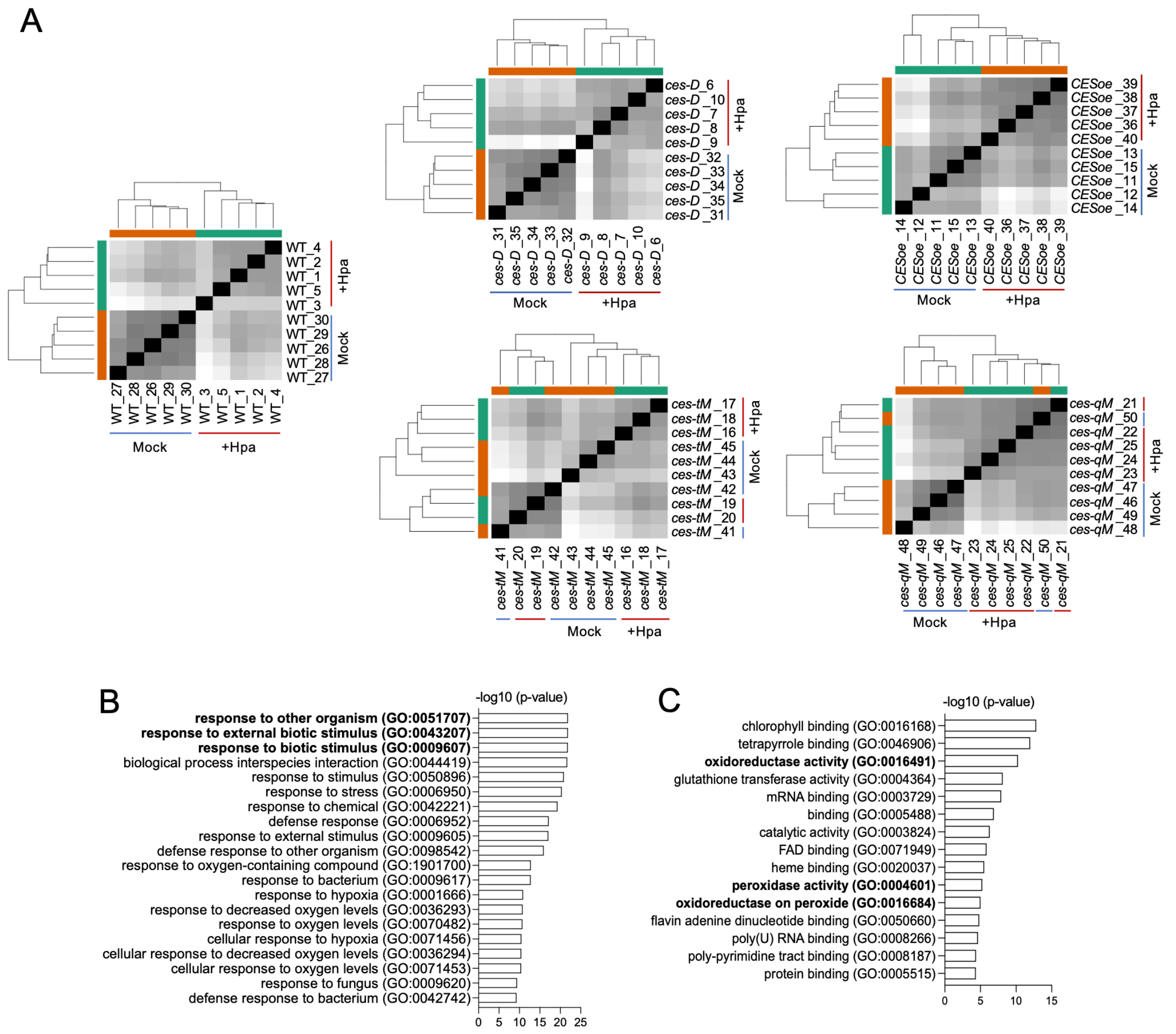


**Fig. S2. RNA-Seq data analysis.**

(*A*) Clustering of RNA-seq data by genotype across treatments. (*B*) Gene Ontology (GO) enrichment of biological processes among upregulated DEGs in cluster 1 of the 193 mutant-specific DEGs. (*C*) GO enrichment of molecular functions among upregulated DEGs in cluster 1 of the 620 DEGs commonly regulated in WT and all *ces* mutants. GO terms are ranked by significance. GO annotations were performed using the PANTHER GO Ontology database with Fisher's exact test and FDR correction ($P < 0.05$).

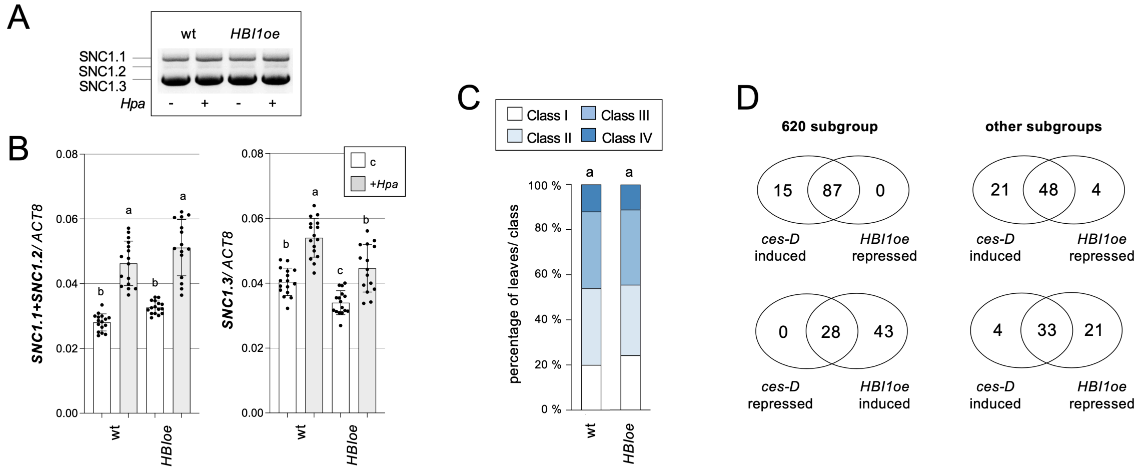


**Fig. S3. Over-expression of *HBI1* neither significantly alters *SNC1* splicing nor *Hpa* Noco2 resistance.**

(*A*) Detection of alternatively spliced *SNC1* transcripts by semi-quantitative PCR in wt and *HBI1oe* plants, either untreated (-) or infected with *Hpa* (+). (*B*) Absolute expression levels of *SNC1.1+SNC1.2* (left) and *SNC1.3* (right) in 18-day-old plants of the *wt and HBI1oe* lines, measured by qPCR. Data represent mean ±SD from four biological replicates, each measured in triplicate and normalized to *ACT8*. (*C*) *Hpa* colonization in the same genotypes as in a. Two-week-old plants were spray-inoculated with *Hpa* Noco2, and colonization was assessed microscopically at 6 dpi in trypan blue-stained leaves. Infection was categorized into four classes; data represent the percentage of leaves per class (n = 200 from 35 plants per genotype). Significance was determined with pairwise Fisher's exact tests. (*D*) Venn diagrams showing overlapping up- and down-regulated DEGs between RNAseq groups containing *ces-D* and *HBI1oe* RNAseq data.

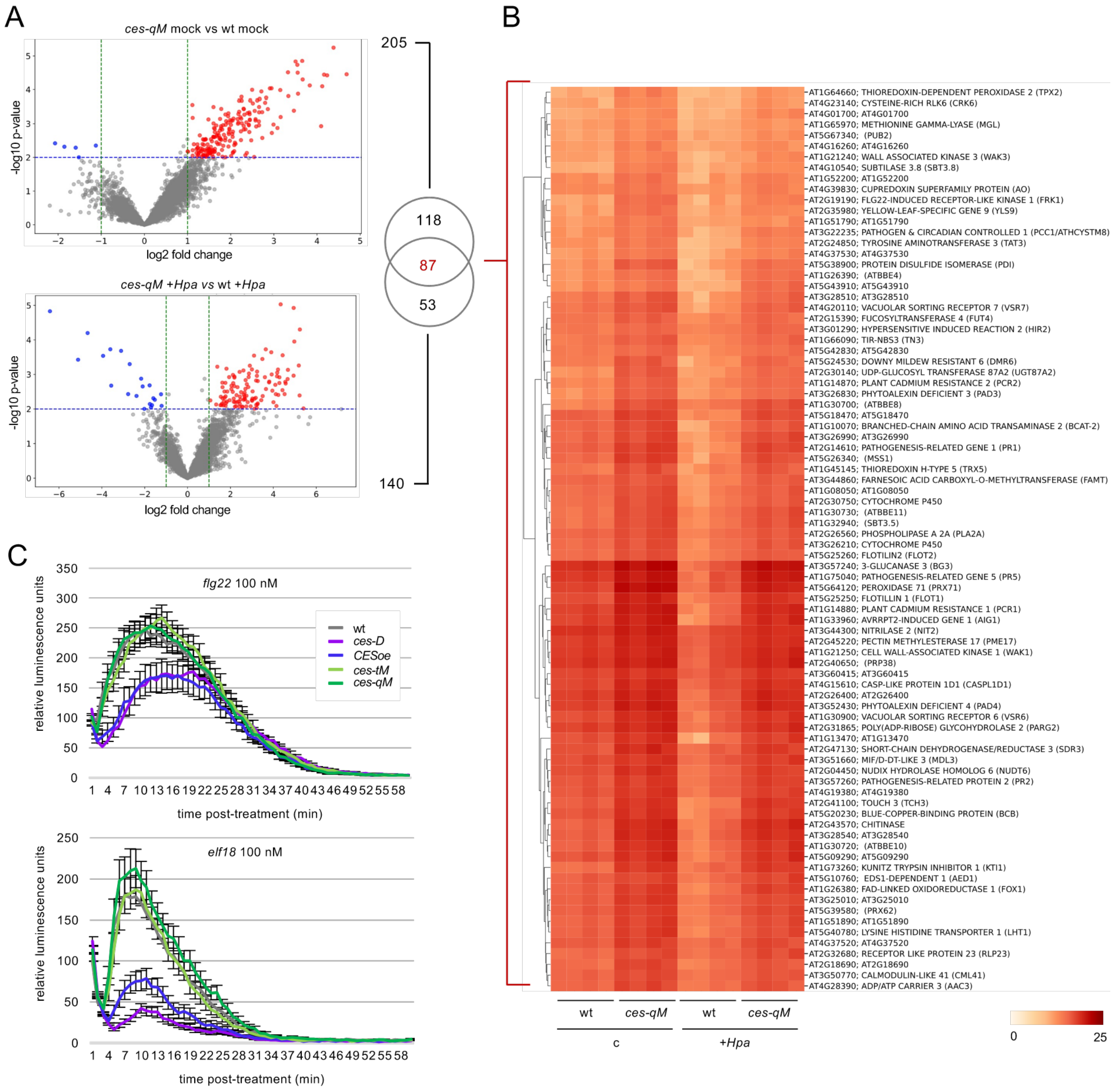


**Fig. S4. CES represses ROS responses.**

(*A*) Volcano plots of significantly differentially abundant proteins in mock-treated *ces-qM* (top) vs wt, and *Hpa*-infected *ces-qM* vs wt at 21 dpi (bottom). Significance was determined using FDR = 0.01 and s0=0.5 thresholds. Overlapping Majority Protein IDs between conditions are shown in the accompanying Venn diagram. (*B*) Heat map of protein abundance for the 84 overlapping Majority Protein IDs. Absolute values were clipped to visualize differential abundance across samples. (*C*) ROS bursts in leaves of wt and *ces* mutant plants following treatment with 100 nM flg22 (top panel) or elf18 (bottom panel). Data represent mean ± SE (n = 8).

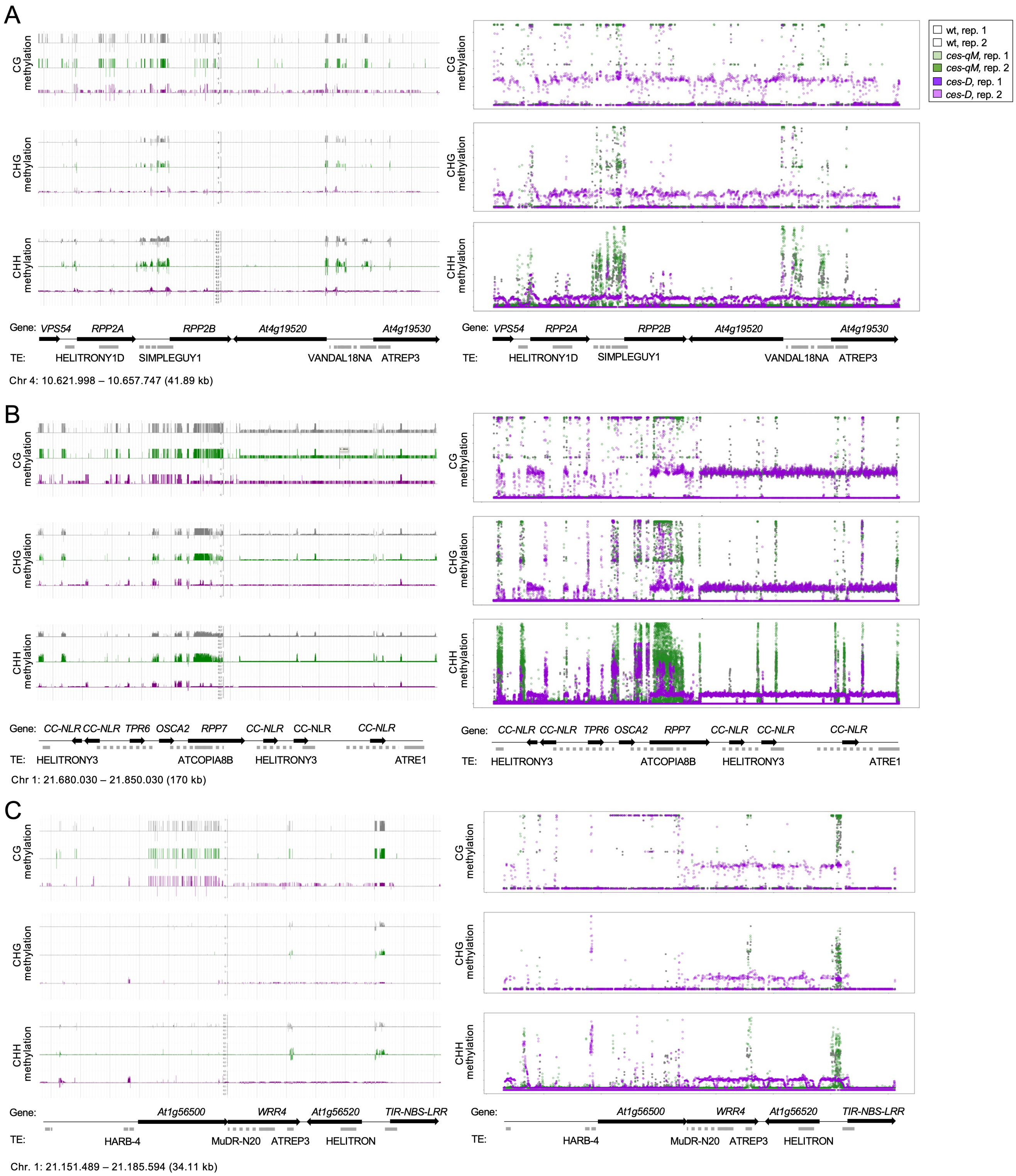


**Fig. S5. CES confer DNA methylation changes in different NLR-containing chromosome region.**

(*A*) *RPP2A/B* cluster. (*B*) *RPP7* cluster. (*C*) *WRR4* cluster. Left: Epigenome browser views (JBrowse) showing plus- and minus-strand methylation in the CG, CHG and CHH context for wt (grey), *ces-qM* (green) and *ces-D* (purple). Right**:** Bubble plots visualizing plus-strand CG, CHG, and CHH methylation in the same chromosome region as on the left, calculated from raw bigWig files using a custom Python script. Y-axes indicate methylation proportion per context; thresholds are auto-scaled per dataset. The location of genes and TEs in these chromosome regions is illustrated below the browser views and plots.

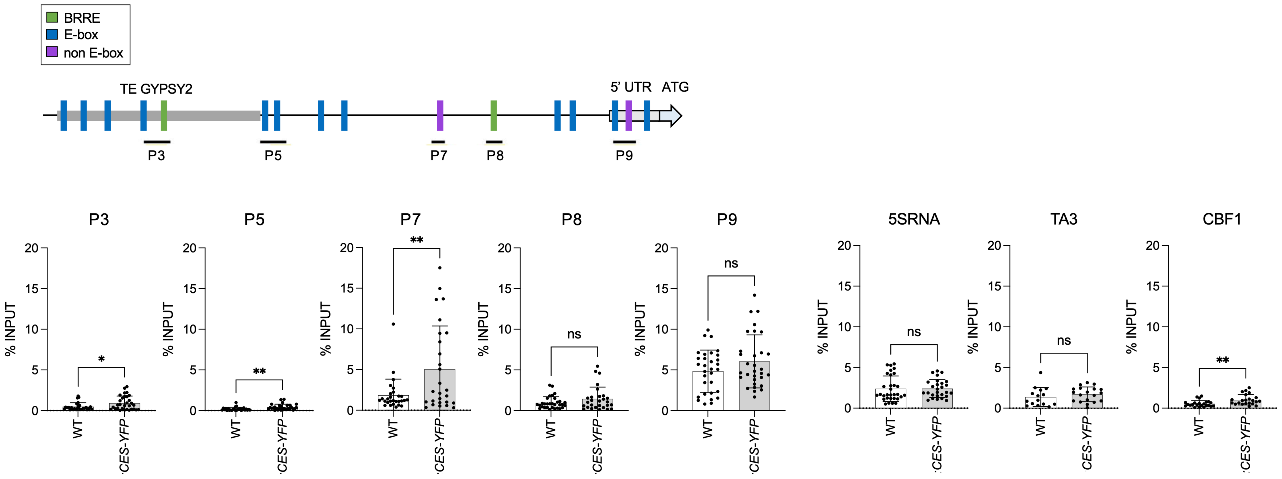


**Fig. S6. CES can bind to the *SNC1* promoter *in vivo*.**

ChIP was performed to assess CES-YFP binding to the *SNC1* promoter. *35S:CES-YFP* and wt plants grown at 21 °C were used and CES-YFP was immunoprecipitated using α-GFP beads. The indicated genomic regions (schematized above) were quantified in the precipitates by qPCR, and enrichment was calculated as the ratio of signal with antibody to without antibody. *5SRNA* and *TA3* served as negative, and *CBF1* as positive controls. Data represent mean ± SE from three biological replicates, each measured in four technical repeats. Bars show SD. Statistical significance was determined by Student's *t*-test (ns P > 0.05, * P ≤ 0.05, ** P ≤ 0.01, *** P ≤ 0.001, **** P ≤ 0.0001).

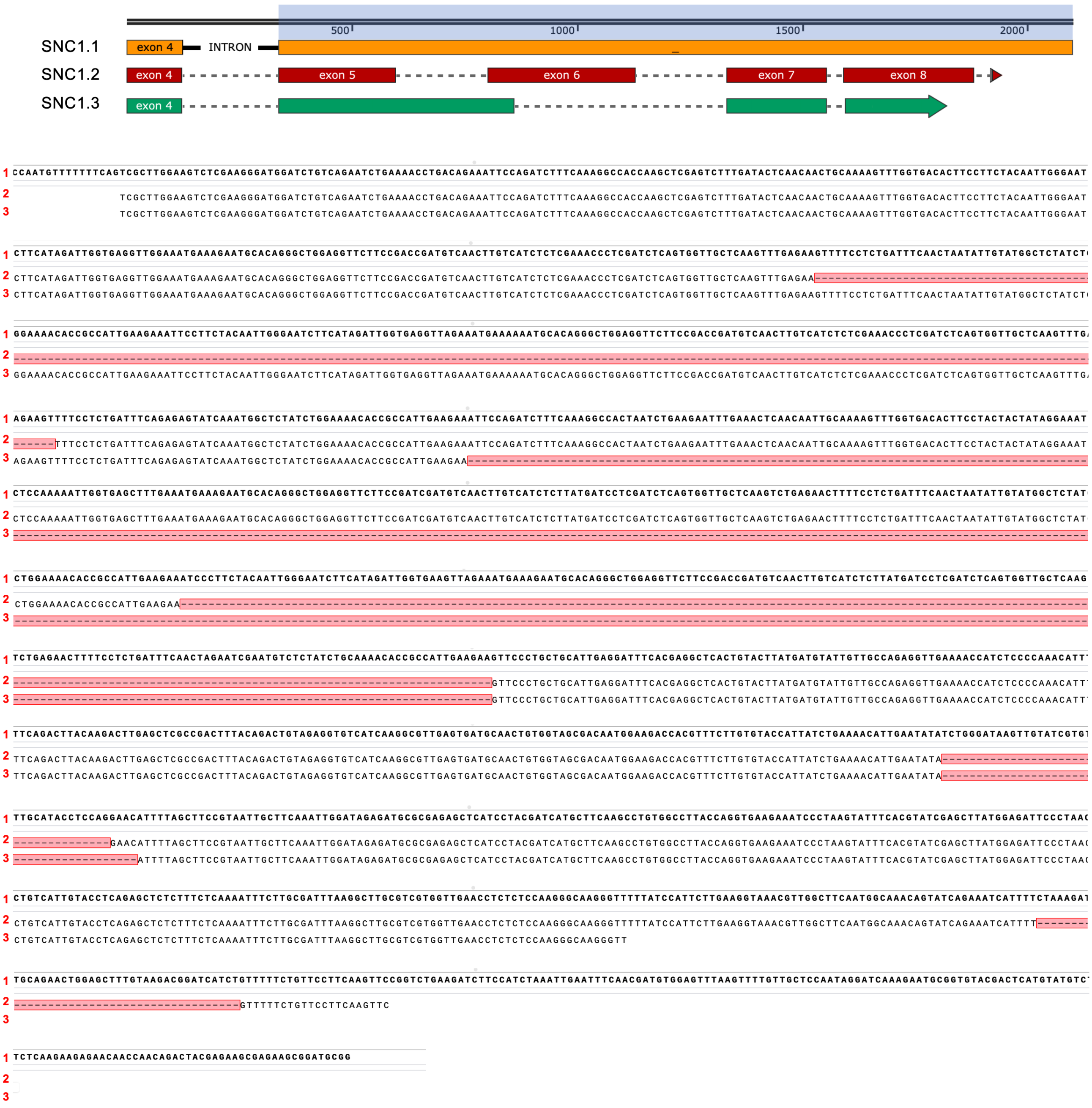


**Fig. S7. The *SNC1* mRNA is alternatively spliced.**

Alignment of genomic sequence of *SNC1* (*At4g16890.1*, TAIR) with sequences obtained from purified PCR-amplified isoform bands derived from cDNA of wt or *ces-D* plants. Sequence 1 corresponds to isoform *SNC1.1*, sequence 2 to *SNC1.2*, and sequence 3 to *SNC1.3*. Schematic representations of these isoforms are shown above the alignment.

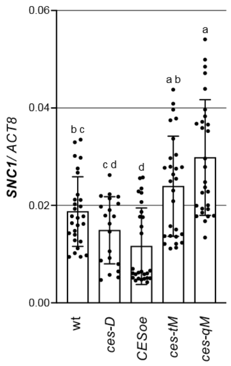


**Fig. S8. Overall *SNC1* transcript abundance in *ces* mutants.**

Absolute expression levels of *SNC1* in 18-day-old plants of *ces* mutants and wt, quantified by qPCRs and using primers that detect a conserved sequence of exon 2 that's present in all splice isoforms. Data represent mean ± SD from four biological replicates, each measured in triplicate and normalized to *ACT8*. Statistical significance was assessed using one-way ANOVA with Tukey's HSD post hoc test in GraphPad Prism v10. All tests were two-sided; $P < 0.05$ was considered significant.

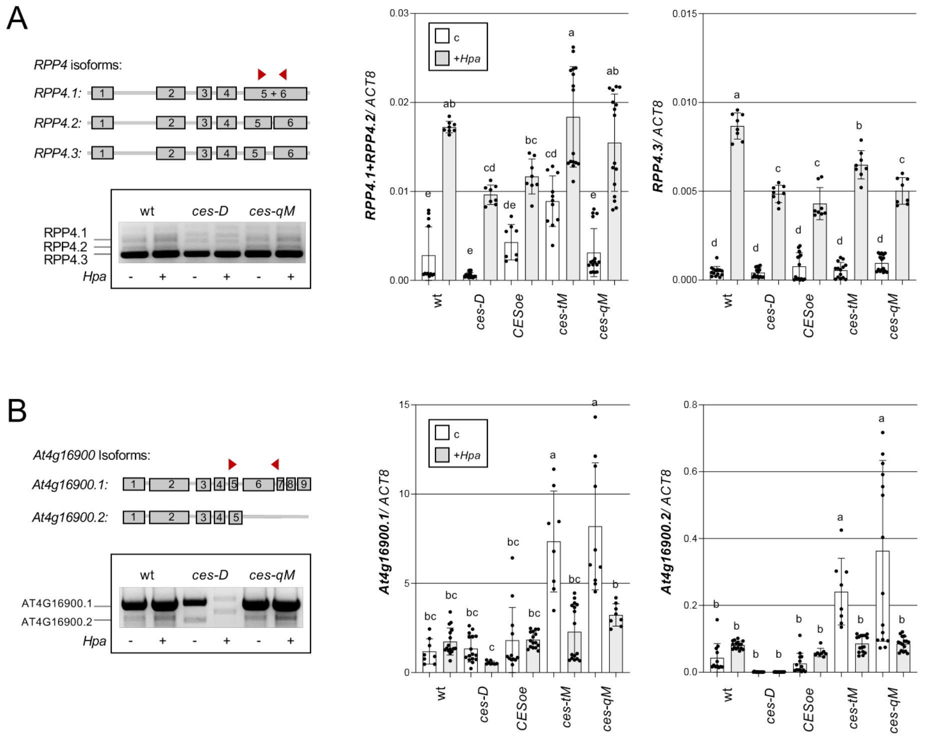


**Fig. S9. CES modulates *RPP4* and *AT4G16900* splicing.**

Detection of predominant *RPP4* (*A*) and *At4g16900* (*B*) mRNA splice variants in wt and *ces* mutants. Left: Scheme of predominant isoforms altered in *ces* mutants with primer positions used to distinguish them indicated by red arrows (top). Detection of alternatively spliced transcripts by semiquantitative PCR in wt, *ces-D*, and *ces-qM* plants, either untreated (-) or infected with *Hpa* (+), using the primers shown (bottom). Right: Absolute expression levels of the indicated isoforms measured by qPCR. Data represent mean ± SD from four biological replicates, each measured in triplicate and normalized to *ACT8*. Statistical significance was assessed using one-way ANOVA with Tukey's HSD post hoc test in GraphPad Prism v10. All tests were two-sided; $P < 0.05$ was considered significant.

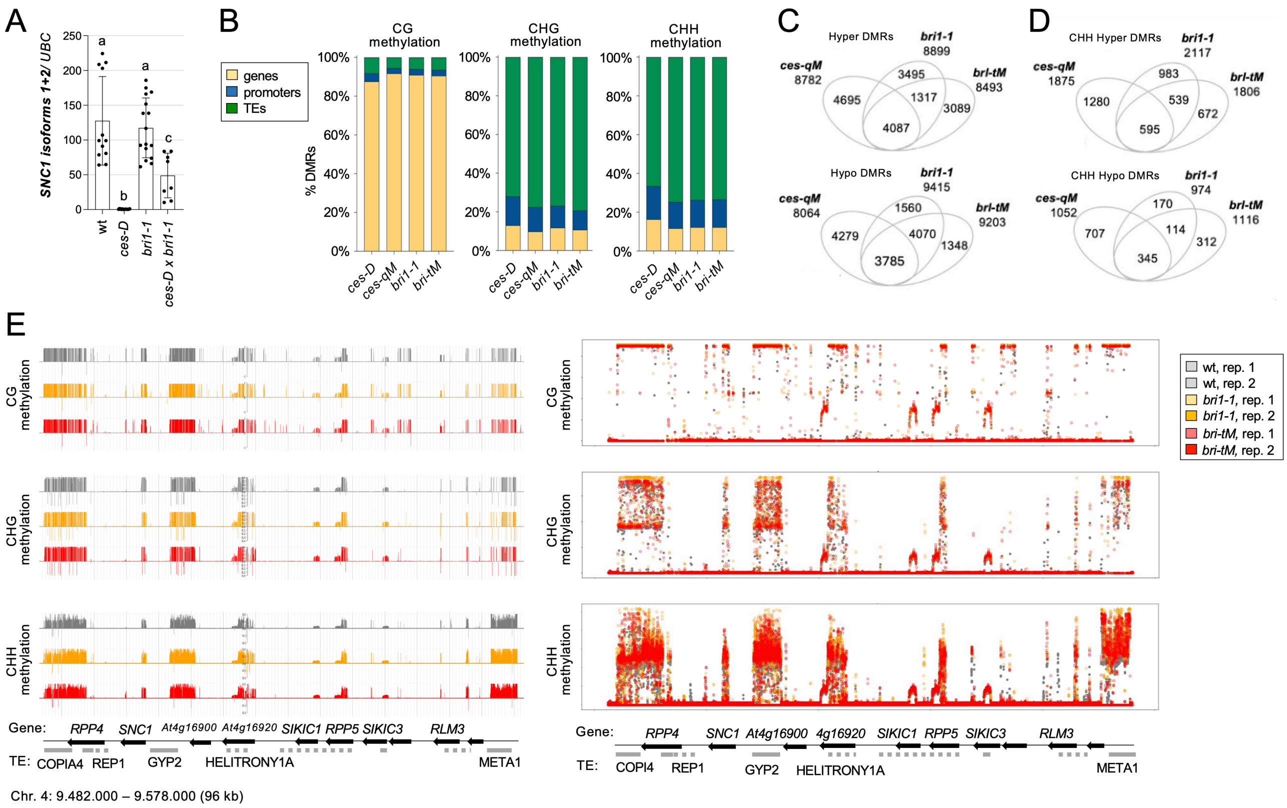

**Fig. S10.** ***BRI1*** **loss-of-function confer** ***ces*** **mutant-like patterns of DNA methylation changes.**

A) Relative expression of *SNC1* in *bri1-1* and *brl-tM* compared to wt and *ces-D.* B) Global distribution of CG, CHG, and CHH-context methylation in *bri1-1* and *brl-tM* mutants, as compared to *ces-D* and *ces-qM*, showing DMR counts identified using METHimpute. C) Venn diagram depicting overlap between hypo- and hyper-methylated DMRs in *brl-tM* and *bri1-1* backgrounds. D) Overlap of CHH-context DMRs (hypo- and hypermethylated) between *brl-tM* and *bri1-1* mutants. E) Visualization of methylation changes at the *RPP5* gene cluster. Epigenome browser views (JBrowse, left) and bubble plots (right) display strand-specific methylation in CG, CHG, and CHH contexts for wt (grey), *bri1-1* (yellow), and *brltM* (orange). Y-axes indicate methylation proportion per context; scales are automatically adjusted per dataset. Gene and TE annotations for the region are shown below.

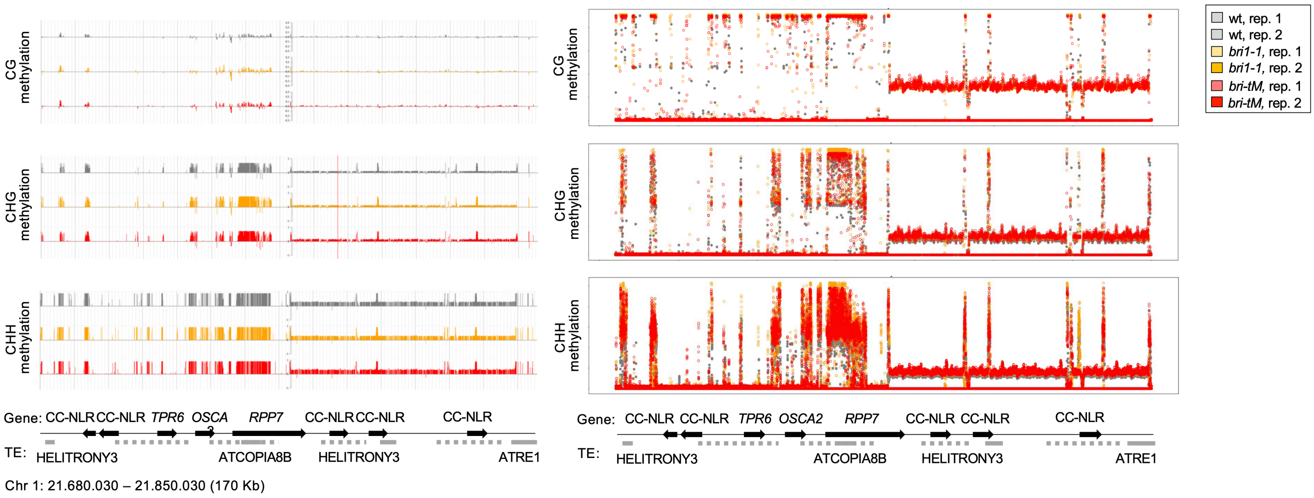


**Fig. S11. *bri1-1* and *brl-tM* mutants show DMRs in TE-containing regions of the *RPP7* locus region.**

Visualization of methylation changes at the *RPP7* gene cluster. Epigenome browser views (JBrowse, left) and bubble plots (right) display strand-specific methylation in CG, CHG, and CHH contexts for wt (grey), *bri1-1* (yellow), and *brl-tM* (orange). Y-axes indicate methylation proportion per context; scales are automatically adjusted per dataset. Gene and TE annotations for the region are shown below.

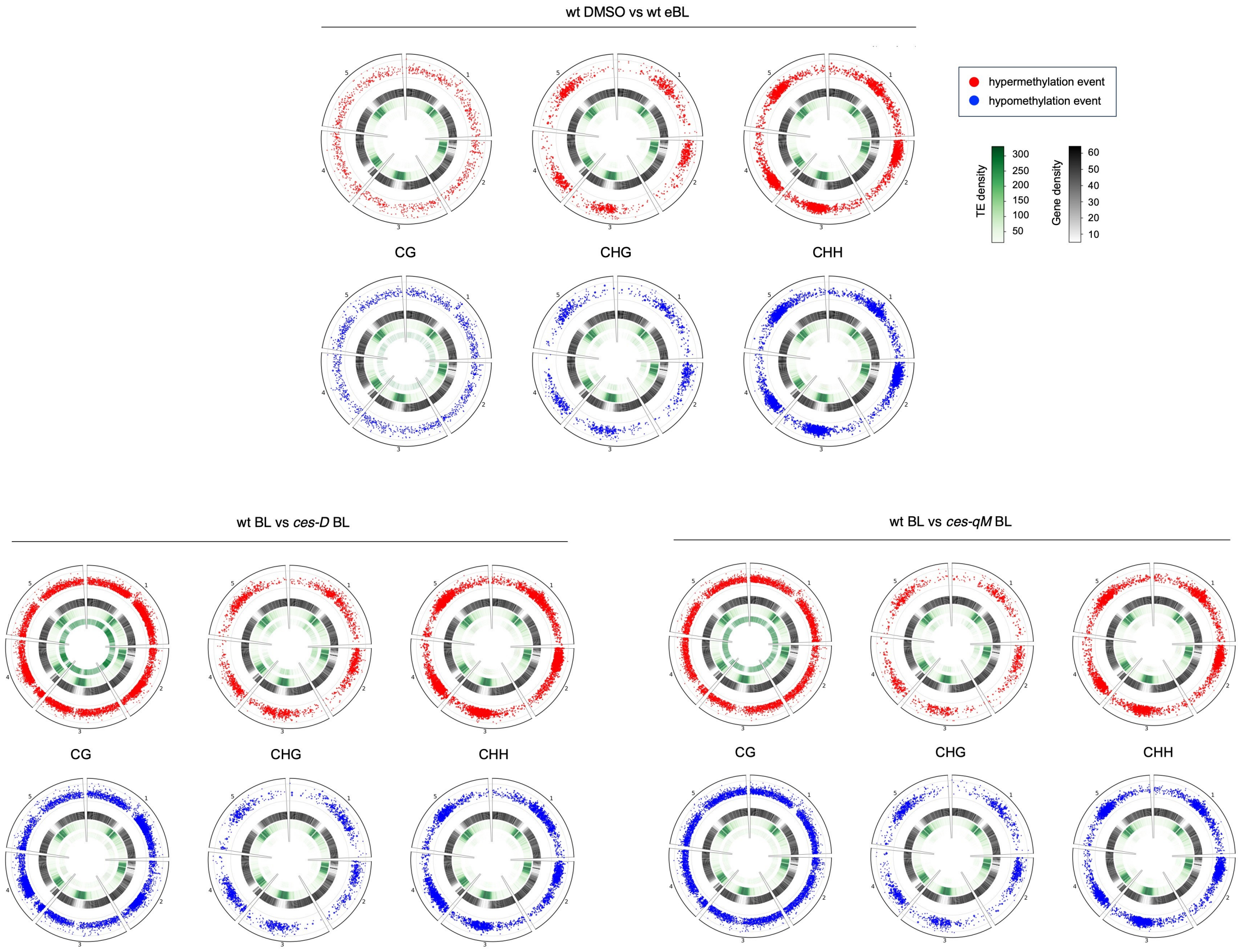


**Fig. S12. BRs induce global DNA methylation changes that are altered in *ces* mutants.**

Global distribution of DMRs in *ces* mutants relative to wt, after epi-BL treatment, categorized by CG, CHG, and CHH sequence contexts, and hypo- or hyper- methylation events. DMRs were identified using METHimpute.

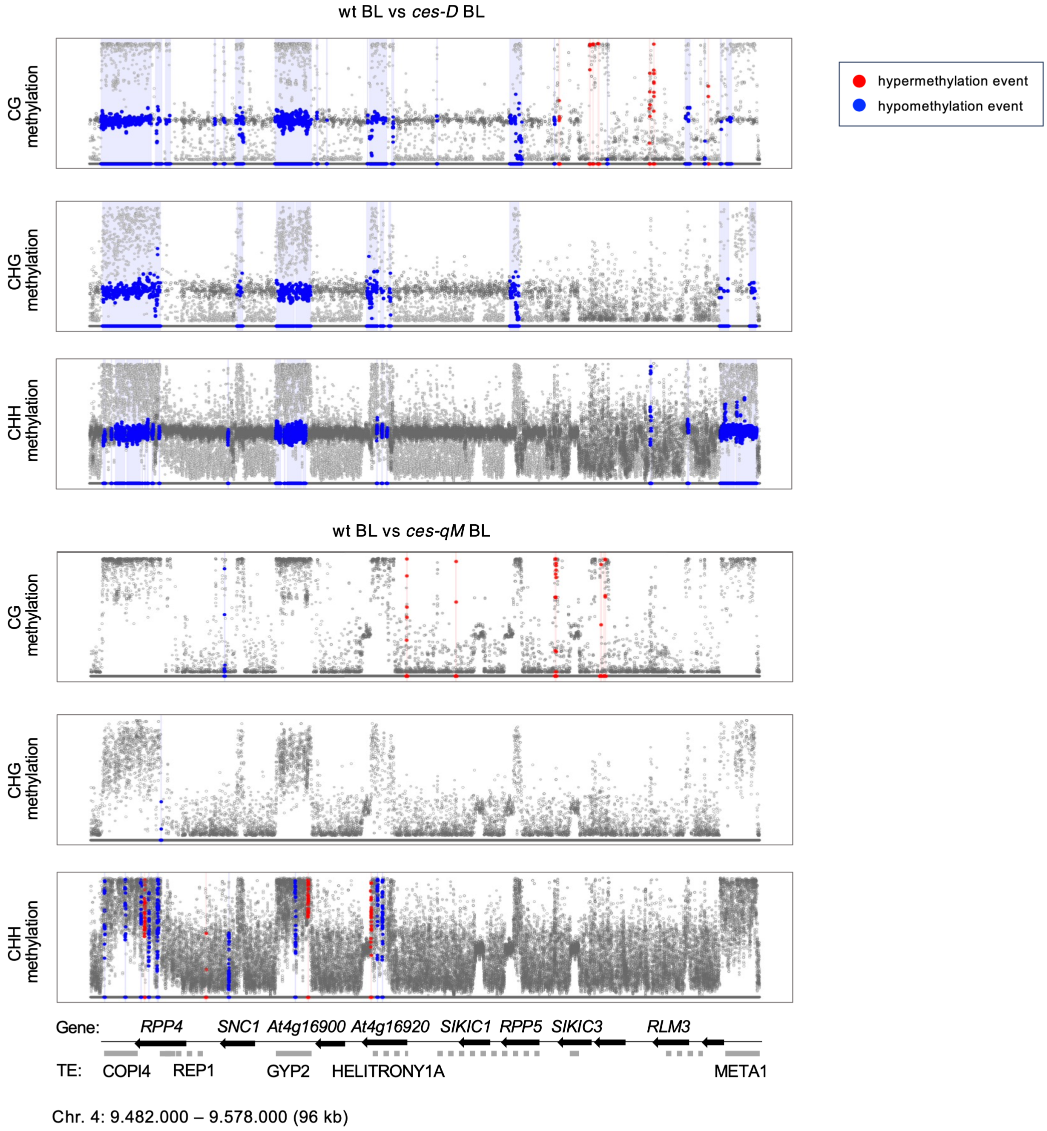


**Fig. S13. CES mutation alters BR-induced DNA methylation changes.**

Visualization of differences in BR-induced methylation changes in the *RPP5* gene cluster region between wild-type and *ces-D* (top) or *ces-qM* (bottom). Bubble plots highlight strand-specific methylation differences after treatment with 24-epiBL, in CG, CHG, and CHH contexts. Red points indicate DMR events called as hyper-methylated, and blue points as hypo-methylated. Y-axes indicate methylation proportion per context; scales are automatically adjusted per dataset. Gene and TE annotations for the region are shown below.

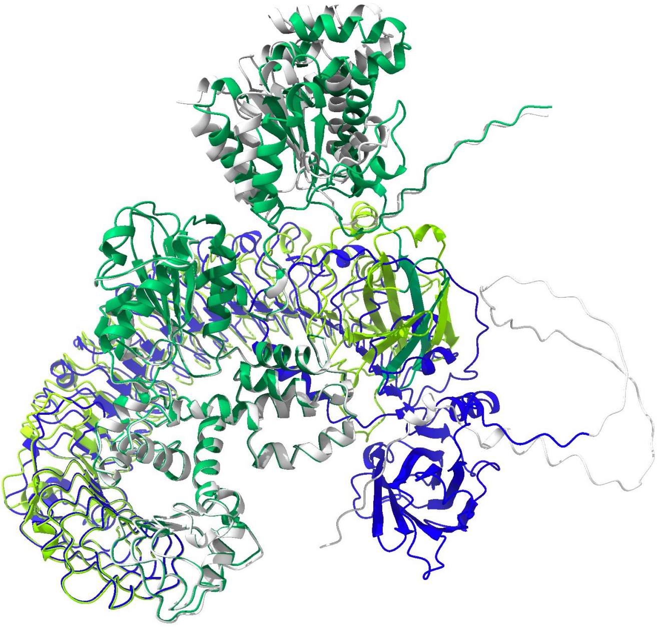

**Fig. S14. AlphaFold3 sequence-based structural modeling of SNC1 isoforms.**

The full-length SNC1.1 protein (grey) is compared with the truncated SNC1.3 isoform (green). Full-length LRR of SNC1.1 is highlighted in blue, and LRR of SNC1.3 in light green. Model built with aligned AA sequences from AlphaFold3 and generated with ChimeraX v1.11
